# Systems analysis uncovers early temozolomide responses and peptide antigens in glioblastoma

**DOI:** 10.1101/2025.11.10.687735

**Authors:** Elizabeth Y. Choe, Owen Leddy, Cecile Riviere-Cazaux, Danielle M. Burgenske, Zeng Hu, Ann C. Mladek, Bogdan I. Fedeles, Sabrina Hu, John M. Essigmann, Robert M. Prins, Jann N. Sarkaria, Terry C. Burns, Rachael A. Vaubel, Forest M. White

**Affiliations:** Koch Institute for Integrative Cancer Research, Massachusetts Institute of Technology, Cambridge, MA; Department of Biological Engineering, Massachusetts Institute of Technology, Cambridge, MA; Ragon Institute of Mass General Brigham, MIT, and Harvard, Cambridge, MA; Department of Neurologic Surgery, Mayo Clinic, Rochester, MN; Department of Radiation Oncology, Mayo Clinic, Rochester, MN; Center for Environmental Health Sciences, Massachusetts Institute of Technology, Cambridge, MA; Department of Chemistry, Massachusetts Institute of Technology, Cambridge, MA; Department of Molecular and Medical Pharmacology, David Geffen School of Medicine at UCLA, Los Angeles, CA; Parker Institute for Cancer Immunotherapy, San Francisco, CA; Department of Laboratory Medicine and Pathology, Mayo Clinic, Rochester, MN

## Abstract

Temozolomide (TMZ) is the standard treatment for nearly all glioblastoma (GBM) patients, as it is the only chemotherapy shown to extend overall survival. However, this benefit is limited to a few months, underscoring the need for combination strategies to improve its efficacy. While TMZ-induced DNA damage can both mediate cytotoxicity and promote resistance, DNA damage more broadly can also stimulate immune activation. To evaluate its immunomodulatory potential, we characterized the previously unexplored early, cell-intrinsic consequences of TMZ in GBM cells, spanning DNA damage, stress responses, and antigen presentation.

A multi-omics approach combining RNA sequencing and quantitative liquid chromatography-tandem mass spectrometry (LC-MS/MS) profiled changes in gene expression, nascent protein translation, steady-state protein levels, kinase-substrate phosphorylation patterns, and MHC-I peptide presentation in GBM cells within 72 hours of TMZ exposure. This analysis revealed rapid activation of DNA damage signaling and p53-associated stress pathways, alongside dynamic changes in protein synthesis and antigen presentation. A set of TMZ treatment-associated peptide antigens (TAPAs) was identified, including peptides derived from stress response proteins, phosphorylated MHC-I peptides, and those induced by other genotoxic treatments such as radiation. Several of these peptides were also detected in recurrent GBM patient tumors.

Our findings suggest that TMZ not only triggers early adaptive and potentially resistance-associated stress programs but may also enhance the immune visibility of GBM cells. These data highlight potential windows for combination therapies with TMZ that bolster immune recognition of GBM, while the systems approach provides a framework to examine how genotoxic therapies across cancers alter tumor immunogenicity.

## Introduction

Temozolomide (TMZ), a DNA-methylating agent, is a frontline treatment for patients with glioblastoma (GBM), the most common and aggressive malignant brain tumor in adults. TMZ induces cytotoxicity by methylating DNA, including at the O^6^ position of guanine, which can lead to double-stranded DNA breaks, cell cycle arrest, and apoptosis.^1^ Although TMZ in combination with surgical resection and concomitant radiation is the standard treatment for GBM, it extends survival by a few months,^2^ with most GBM patients surviving less than two years post-diagnosis. Moreover, over half of GBM tumors exhibit intrinsic or acquired resistance to TMZ.^3^ As such, opportunities remain to enhance the therapeutic efficacy of TMZ in GBM patients.

Paradoxically, while DNA damage can drive therapeutic resistance, it may also create new therapeutic opportunities. DNA damage and immune activation are closely linked across a variety of physiological processes, including infectious diseases, aging, and cancer.^4,5^ The resulting signaling cascades from DNA damage can lead to cytokine and interferon release, which activate and recruit immune cells to target abnormal cells. In several cancers, DNA-damaging therapies such as radiation and certain chemotherapies have increased the surface presentation of major histocompatibility complex class I (MHC-I) molecules, which present cell-derived peptides to the immune system.^6–9^ Additionally, recent studies have identified therapeutically-targetable, treatment-associated peptide antigens (TAPAs),^10–12^ which are peptides upregulated on MHC-I molecules by therapies like kinase inhibitors, that create potential drug-inducible tumor vulnerabilities.

While the long-term response to TMZ has been characterized in recurrent GBM,^1,13^ less is known about the GBM cellular response that occurs within the first few days of TMZ treatment, particularly with regard to TAPAs. Understanding the dynamics of MHC-I presentation shortly after TMZ exposure may reveal novel targets or therapeutic windows before immune evasion occurs. This early characterization may inform optimal timing to maximize efficacy of immunotherapies such as immune checkpoint inhibitors (ICIs), bispecific T cell engagers (BiTEs), T cell engagers (TCEs), or peptide-targeting vaccines, which require sufficient presentation of tumor-specific or tumor-associated peptide-MHC complexes. This is particularly relevant in GBM’s immunologically cold environment, where strategies to improve immunotherapy response remain under investigation.^14,15^

Traditional sequencing approaches have historically provided only a partial view of these dynamics. While GBM tumor cell response to TMZ has been genomically and transcriptomically characterized, these data only partially correlate with protein expression or TAPAs, which are often the direct targets of therapeutic interventions.^16,17^ Moreover, technical barriers have historically limited large-scale, systems-level characterization of these dimensions of cellular regulation. However, advances in quantitative mass spectrometry now enable profiling of proteomes and phosphoproteomes, as well as newly translated proteins and immunopeptidomes, all with high resolution.^10,18,19^

Here, we applied a multi-omics approach to capture a broad spectrum of previously unexplored early dynamic cellular responses to TMZ in human GBM cells. We sought to understand the multifactorial impact of TMZ and identify opportunities to improve its efficacy. This approach combined RNA sequencing with recent liquid chromatography-tandem mass spectrometry (LC-MS/MS) approaches in quantitative translatomics, proteomics, phosphoproteomics, and immunopeptidomics to directly profile early changes in mRNA, nascent protein synthesis, global protein abundance, phosphoproteomic signaling, and the repertoire of MHC-I peptides in TMZ-treated GBM cells. Our analysis revealed rapid activation of DNA damage signaling and p53-driven stress responses, along with dynamic changes in translation and antigen presentation. We further characterized, for the first time, TMZ-induced TAPAs, several of which were also detected in recurrent GBM patient tumors and in cells treated with other genotoxic therapies. These results provide a systems-level view of the cell-intrinsic consequences of TMZ-induced genotoxic stress and highlight potential combinations that could improve GBM therapy.

## Materials and Methods

(LC/MS analysis and peptide identification and quantitation methods available in Supplementary Methods)

### Cell culture and drug treatment

U87-MG cells were obtained from ATCC (ATCC HTB-14^TM^; RRID:CVCL_0022); GBM177 and GBM181 patient-derived cell lines were provided by the Mayo Clinic Brain Tumor Patient-Derived Xenograft National Resource. All cell lines were maintained in Dulbecco’s modified Eagle’s medium (DMEM; Corning) supplemented with 10% fetal bovine serum (FBS; Gibco) at 37°C, 5% CO_2_. Cells were passaged at a 1:4 ratio, passaging no more than three times from a thawed vial, and were routinely tested for mycoplasma using a detection kit (Lonza). Cell line identity was confirmed by the source repository, and no further authentication was performed by the authors.

For TMZ vs. DMSO experiments, three replicates were generated for each timepoint/condition by thawing separate vials of cells and expanding them into 10 cm dishes. Cells were seeded at a density of 10^6^ cells per 10 cm dish and allowed to adhere for 24 hours before replacing media with fresh media containing either 100 μM TMZ (SelleckChem) or 0.1% dimethyl sulfoxide (DMSO; Sigma) as vehicle control. Multiple plates per replicate were treated in parallel to provide sufficient material for omics workflows. Because these were clonal cell populations expanded in culture, randomization of plates was not applicable, and investigators were aware of treatment conditions.

For immunopeptidome studies comparing various genotoxic therapies, cells were treated with low and high doses of the following: TMZ (10 µM and 900 µM; SelleckChem), BI 907828 (Brigimadlin; 10 nM and 1 µM, provided by the Sarkaria lab), radiation (2.5 Gy and 9 Gy; Gammacell 40 Exactor, Theratronics), and interferon-gamma (IFN-γ; 10 ng/mL and 100 ng/mL; Thermo Fisher). Each condition was run as a single sample to allow direct comparison of high-vs. low-dose treatments across all five conditions within the constraints of TMT channel capacity.

### Patient GBM tumor samples

First-recurrence GBM tumors were obtained from the Mayo Clinic in Rochester, Minnesota and multiple-recurrence tumors from the David Geffen School of Medicine at UCLA/LA BioMed at Harbor-UCLA Medical Center.

At the Mayo Clinic, patients underwent surgical resection of their GBM at the time of suspected first recurrence per standard neurosurgical protocol. Resected tissue was flash-frozen and stored under the Mayo Clinic Neuro-Oncology Biobank (IRB#12-003458). All participants provided written informed consent, and tissue analysis for this study was approved under protocol IRB#17-004675. This study was performed in accordance with the Declaration of Helsinki and approved by the Mayo Clinic Institutional Review Board.

At UCLA, surgical resection of tumor tissue of GBM patients with recurrent patients with multiply recurrent GBM were treated with off-label neoadjuvant immunotherapy immediately prior to surgical resection. Tumor tissue was resected, flash-frozen, and stored under the UCLA institutional tumor bank protocol (IRB#10-0065). All participants provided written informed consent, and the study was conducted in accordance with the Declaration of Helsinki.

All available recurrent GBM tumor specimens provided by collaborators were considered. Patient samples were included if they had suspected recurrent GBM and if there was sufficient tumor weight after surgery for downstream omics analyses. Samples were excluded if tumor material was insufficient. No additional exclusions were made. Both male and female patients were included, and demographic and clinical information (age, sex, recurrence number, and prior treatments) is provided in Supplementary Table S2.

### Flow cytometry

After 24, 48, and 72 hours exposure to 100 μM TMZ (SelleckChem), 0.1% DMSO (Sigma), or 50 ng/mL INF-γ (Thermo Fisher), cells were lifted with 0.05% Trypsin-EDTA (Thermo Fisher) for 5 minutes, spun at 300 x *g* for 5 minutes at 4°C, and washed with cold flow buffer made of 1% bovine serum albumin (BSA; Sigma) in phosphate-buffered saline (PBS; Corning). Cells were stained with 3 μg/mL Alexa Fluor 488® HLA-A, B, C antibody (BioLegend, RRID:AB_493134) in flow buffer for 30 minutes on ice in the dark, then rinsed twice with flow buffer. Cells were resuspended in flow buffer with 2 μg/mL DAPI staining solution (Thermo Fisher, RRID:AB_2629482). Samples were analyzed using a FACS Celesta HTA-1 (BD Biosciences) and FlowJo software version 10.10.0 (RRID:SCR_008520).

### GDF15 ELISA

After 24, 48, and 72 hours, cell media was frozen at -80°C. Media was diluted 1:5 or 1:50 and GDF15 concentration was measured using a Human GDF15 ELISA kit (Thermo Fisher), according to the manufacturer’s instructions.

### RNA sequencing

Total RNA was isolated from samples after 24, 48, and 72 hours of TMZ or DMSO treatment using the PureLink^TM^ RNA Mini Kit (Invitrogen) according to the manufacturer’s instructions. RNA was cleaned using RNA-SPRI beads (Aline Biosciences), and quality and concentration were assessed using a Fragment Analyzer (Agilent Technologies). For each sample, 500 ng of RNA was used for rRNA depletion with the NEBNext rRNA Depletion Kit, followed by library preparation using the NEBNext Ultra II Directional RNA Library Prep Kit for Illumina (New England Biolabs). Libraries were sequenced on an Illumina NextSeq500 with 40 nt paired-end reads.

RNA-seq data were processed using nf-core/rnaseq v3.12.0,^20^ executed with Nextflow v22.10.4 (RRID:SCR_024135),^21^ using the --aligner star_salmon and --pseudo_aligner salmon options. Reads were aligned to the GRCh38 primary assembly using STAR (RRID:SCR_004463), and transcript abundance was estimated using Salmon (RRID:SCR_017036) with the GENCODE v43 annotation (RRID:SCR_014966). Processed RNA-seq results are provided in Supplementary Table S3.

### Sample preparation for TMT-labeled proteome and phosphoproteome analysis

Samples were prepared adapting a previously described protocol.^18^ After 24, 48, and 72 hours of exposure to TMZ or DMSO, cell plates were moved to ice, rinsed with cold PBS, and lysed with 500 μL 8M urea and cell scraping. Lysates were cleared by centrifugation at 3,500 × *g* for 5 minutes at 4°C, followed by bicinchoninic acid assay (BCA; Pierce) to determine protein concentration. Roughly 1 mg of proteins were reduced with 10 mM dithiothreitol (DTT) for 30 minutes at 56°C, then alkylated with 55 mM iodoacetamide for 1 hour at room temperature, rotating in the dark. Chemical modifications were quenched and urea diluted by adding 4× the reaction volume of 100 mM ammonium acetate pH 8.9. Digestion took place for 18 hours, rotating at room temperature, with sequencing grade modified trypsin (Promega) at a 1:50 ratio of trypsin to total protein. The digest was quenched by adding 10% of the total reaction volume of glacial acetic acid. Peptides were then desalted using Sep-Pak Plus Light C18 solid-phase extraction cartridges (Waters). Peptides were eluted from the cartridges with 7 mL of 40% acetonitrile in 0.1% acetic acid and dried to approximately 2 mL using vacuum centrifugation. Peptide concentration was then measured by BCA and lyophilized in 150 μg aliquots.

Peptides were resuspended in 35 μL 50 mM pH 8.5 HEPES. TMTpro 16-plex (490 μg; Thermo Fisher) was resuspended in 15 μL anhydrous acetonitrile and added to each sample; peptides were labeled for 5 hours at room temperature on a shaker. Labeling reactions were quenched using 0.3% hydroxylamine, pooled, with sample tubes rinsed twice with 20 μL 25% acetonitrile in 0.1% acetic acid and dried by vacuum centrifugation overnight. Dried, labeled peptides were stored at -80°C until analysis.

Serial immunoprecipitation (IP) enabled motif-specific phosphoproteomics and total protein analysis of the same samples. First, phospho-SQ/TQ containing peptides were enriched using the PTMScan® Phospho-ATM/ATR Substrate Motif Kit (Cell Signaling Technology). Dried peptides were resuspended in 400 μL of immunoprecipitation buffer (100 mM Tris-HCl, 0.3% NP-40, pH 7.4) and added to 38.5 μL of antibody-conjugated beads from the kit, rotating for 18 hours at 4°C. Beads were rinsed once with IP buffer and three times with 100 mM Tris-HCl (pH 7.4), and peptides were eluted twice with 25 μL 0.2% trifluoroacetic acid (TFA). To minimize non-specific binding, all immunoprecipitated phosphopeptides were further enriched using the High-Select™ Fe-NTA Phosphopeptide Enrichment Kit (Thermo Fisher), modifying the manufacturer’s protocol to use 50 μL input peptide and 2 × 20 μL elution volume. Eluates were vacuum-concentrated to <5 μL and resuspended in 10 μL 5% acetonitrile in 0.1% formic acid for LC-MS/MS analysis. The supernatant from the phospho-SQ/TQ IP was subjected to a second IP using the PTMScan® Phospho-MAPK/CDK Substrate Motif Kit (PXS*P and S*PXK/R), with the same subsequent protocol.

The supernatant from the second IP was used for global phosphopeptide enrichment and total protein expression profiling via high-pH reverse-phase fractionation. Half of the supernatant from the second IP was fractionated via high-pH reversed-phase chromatography on a C18 HPLC column (Kromasil) using 10 mM triethylammonium bicarbonate (TEAB) and acetonitrile buffers (Agilent 1100 HPLC). Ten fractions were collected and dried to ∼1 mL by vacuum centrifugation. For each fraction, 90% was reserved for global phosphorylation analysis and 10% for protein expression profiling. Global phosphorylation fractions were enriched using the Fe-NTA Phosphopeptide Enrichment Kit and processed as above. Protein expression fractions were resuspended in 15 μL 5% acetonitrile in 0.1% formic acid. For both global phospho-and protein fractions, 2 μL was injected per LC-MS/MS run.

### Sample preparation for TMT-labeled translatome analysis

Samples were prepared for translatomics adapting a previously-described protocol.^22^ Following 24, 48, or 72 hours of TMZ or DMSO treatment, media was replaced with lysine/arginine/methionine (KRM)-free DMEM (AthenaES) for a 30-minute starvation, followed by supplementation with 3 mM L-azidohomoalanine (AHA) (VectorLabs), 0.5 mM ^15^N_4_^13^C_6_ arginine (R10) (Thermo Fisher), and 0.5 mM ^15^N_2_^13^C_6_ lysine (K8) (Thermo Fisher) to allow for labeling of newly-translated proteins for 1 hour. Cells were washed with ice-cold PBS and lysed via cell scraping in 250 μL of 1% SDS in PBS containing 300 μg/mL cycloheximide, 50 mM N-ethylmaleimide (NEM), and 1:1000 benzonase. Lysates were normalized by BCA assay to 200-500 μg of protein in 500 μL total volume of 1% SDS. An equal volume of 8 M urea with 850 mM NaCl was added.

To control for sample preparation variability, AHA-containing synthetic heavy-isotope–labeled peptides (AHA-SHIPs) were spiked into each lysate prior to enrichment. AHA-SHIPs were synthesized at the MIT Biopolymers and Proteomics Lab using AHA along with ^13^C arginine, ^13^C lysine, or ^13^C/^15^N isoleucine. Three distinct peptides were added to each sample: AHA-GGGGRAP(*I)(*I)AVT(*R), AHA-GGGGRGALQN(*I)(*I)PASTGAA(*K), and AHA-GGGGRGLGTDEDTL(*I)E(*I)LAS(*R). Peptides were stored as 1 mM stocks in 0.1% TFA at -20 °C.

Newly translated proteins were enriched by click chemistry using 30 μL DBCO-agarose beads (VectorLabs) per sample, prewashed 3× with 0.8% SDS in PBS. In addition to the sample lysates, 3 pmol each of the AHA-SHIPs were added to the beads for incubation. Enrichment took place for at least 16 hours at room temperature with rotation. After enrichment, samples were transferred to Micro Bio-Spin Columns (Bio-Rad) to drain. Columns were washed with 1 mL ultrapure water, followed by on-bead reduction using 1 mL of 10 mM DTT in 0.8% SDS in PBS at 50 °C for 30 minutes. Proteins were alkylated with 1 mL 50 mM NEM in 0.8% SDS in PBS at room temperature for 30 minutes. Columns were drained and sequentially washed 8× each with 0.8% SDS in PBS, 8 M urea, and 20% acetonitrile. After the second wash of each set, the columns were capped and allowed to incubate for 10 minutes with the wash buffer before continuing with the remaining washing. After final drainage and drying via quick spin, beads were resuspended in 300 μL digest buffer (200 mM TEAB in 10% acetonitrile) and transferred to a microcentrifuge tube. The column was rinsed twice with 300 μL digest buffer, with rinses pooled into the same microcentrifuge tube. Beads were pelleted (5,000 × g, 5 min), resuspended in 100 μL of digest buffer containing 1 ng/μL trypsin, and digested for 18 hours at room temperature with rotation.

TMTpro 16-plex reagents (490 μg in 15 μL anhydrous acetonitrile) were added directly to each digest and incubated for 5 hours at room temperature on a shaker. Labeling reactions were quenched using 0.3% hydroxylamine, pooled, with sample tubes rinsed twice with 20 μL 50% acetonitrile in 0.1% acetic acid and dried by vacuum centrifugation overnight.

Dried samples, including beads, were resuspended in 400 μL 5% acetonitrile in 10 mM TEAB (pH 8). Samples were loaded onto custom-packed C18 fractionation columns (200 μm inner diameter fused silica capillary packed with 20 cm of 10 μm C18 beads). After loading the sample, columns were rinsed with at least 50 μL of 5% acetonitrile in 10 mM TEAB (pH 8). Peptides were fractionated into 8 parts by high-pH reverse-phase chromatography on an Agilent 1100 Series HPLC using 10 mM TEAB and acetonitrile buffers. Fractions were lyophilized and stored at -80 °C until LC-MS/MS analysis. Before analysis, fractions were resuspended in 10 μL 5% acetonitrile in 0.1% formic acid, with 4 μL per fraction injected for LC-MS/MS.

### Sample preparation for TMT-labeled immunopeptidome analysis

Cell line samples were prepared for immunopeptidomics adapting a previously described protocol.^10^ After 24, 48, and 72 hours, cells were lifted with 0.05% Trypsin-EDTA for 5 minutes, spun at 500 × *g* for 5 minutes at 4°C, washed with ice-cold PBS, and again pelleted. Cells were resuspended in 750 μL-1 mL lysis buffer (20 nM Tris-HCl pH 8, 150 mM NaCl, 0.2 mM phenylmethylsufonyl fluoride (PMSF) (Sigma), 1% CHAPS (Thermo Fisher), and 1X HALT Protease/Phosphatase Inhibitor Cocktail (Thermo Fisher). Cell membranes were disrupted via sonication (5 × 10 seconds microtip sonicator pulses). Lysates were cleared via centrifugation at 5,000 × *g* for 5 minutes at 4°C and protein concentration was determined using BCA.

For each sample, pan-HLA antibody-conjugated beads were prepared for IP by adding 0.5 mg of anti-human MHC class I (HLA-A, HLA-B, HLA-C) clone W6/32 antibody (BioXCell, RRID:AB_1107730) to 20 μL FastFlow Protein A Sepharose bead slurry (GE Healthcare) for 4 hours rotating at 4°C. Beads were washed with 500 μL of an IP buffer (20 nM Tris-HCl pH 8 containing 150 mM NaCl).

To control for sample preparation variability, heavy isotope-coded peptide-MHC standard complexes (hipMHCs) were spiked into each lysate prior to enrichment. Three different heavy isotope-coded peptides (hips) were synthesized at the MIT Biopolymers and Proteomics Lab using ^13^C/^15^N leucine as previously described:^10^ A(*L)ADGVQKV, ALNEQIAR(*L), and SLPEEIGH(*L); hips were stored in 500 μM stocks in 0.1% TFA at -20°C. To create hipMHC complexes, 34 μL PBS, 4 μL of stock hip, and 2 μL recombinant, biotinylated Flex-T HLA-A*02:01 monomers (BioLegend) were combined in a 96-well V-bottom plate. Samples were exposed to UV light (365 nm) for 30 minutes on ice. The concentrations of the hipMHC complexes were determined using the Flex-T HLA class I ELISA kit (BioLegend) according to the manufacturer’s instructions.

MHC-I peptides were enriched by adding equal amounts of protein from each sample (2-8 mg) and 30 fmol each of the three hipMHCs to conjugated beads. The IP took place rotating at 4°C overnight. Beads were then washed once with 500 μL cold tris buffered saline (TBS) (Corning) and twice with 500 μL ultrapure water. MHC peptides were eluted from the beads by adding 500 μL 10% acetic acid to the beads and rotating for 20 minutes at room temperature. The elutions were added to 10K molecular weight cutoff filter columns (PALL Life Science), which had been passivated with 1% bovine serum albumin in PBS for 20 minutes-1 hour, to isolate peptides. The filter column was centrifuged at 1,250 × *g* for 28 minutes and the sample was lyophilized. Lyophilized peptides were resuspended in 33 μL 150 mM TEAB in 50% ethanol. TMTpro 16-plex (80 μg) was resuspended in 10 μL anhydrous acetonitrile and added to each sample; peptides were labeled for 5 hours at room temperature on a shaker. Labeling reactions were quenched using 0.3% hydroxylamine, pooled, with sample tubes rinsed twice with 20 μL 25% acetonitrile in 0.1% acetic acid and dried by vacuum centrifugation overnight. Dried, labeled peptides were stored at -80°C until LC-MS/MS analysis.

Excess TMT was cleaned from samples using single-pot solid-phase-enhanced sample preparation (SP3). First, hydrophobic and hydrophilic Sera-mag carboxylate-modified speed beads (GE Healthcare) were combined in a 1:1 mix with beads at a final concentration of 10 μg/μL in ultrapure water. TMT-labeled samples were resuspended in 30 μL 100 mM ammonium bicarbonate pH 7-8 and added to 500 μg of the bead mixture with 1 mL acetonitrile, shaking for 8 minutes at room temperature. Beads were rinsed twice with 200 μL acetonitrile and peptides were eluted from the beads in 25 μL 2% DMSO, shaking for 5 minutes at room temperature. The elution was dried in a new microcentrifuge tube in a vacuum centrifuge. Peptides were resuspended in 20 μL 3% acetonitrile in 0.1% formic acid and 4 μL was injected for LC-MS/MS analysis.

### Sample preparation for targeted, SureQuant TAPAs analysis

Custom synthetic stable isotope-labeled (SIL) standards to quantify 10 TMZ-associated MHC-I peptides of interest were synthesized using the same approach as the previously described hips, incorporating ^13^C leucine, ^13^C/^15^N glutamic acid, or ^13^C lysine. The peptide sequences were: RQIpSQDV(*K)L (phosphorylated AMPD2), A(*L)WTVQEA (CSF3), A(*L)PEGLPEA (GDF15), D(*E)DPATYVW (HAS1), L(*L)DVPTAAV (IFI30), E(*E)LGFRPEY (PARP1), S(*L)AEVLQQL (RETSAT), S(*E)IAVGHQY (SLC25A35), SIDpSPQ(*K)L (phosphorylated TP53BP1), L(*E)GNPDTHSW (UBAP2L). Peptides were stored in 10 mM stocks in 20% acetonitrile in 0.1% formic acid at -20°C. NetMHCpan-4.1 was used to predict the HLA allele to which each peptide would most likely bind.^23^ To create SIL-MHC complexes, easYmer© kits (ImmunAware) for HLA-A*02:01, HLA-B*44:02, or HLA-C*05:01 were used according to the manufacturer’s instructions. The concentrations of the SIL-MHC complexes were determined using the Flex-T HLA class I ELISA kit (BioLegend), according to the manufacturer’s instructions, and complexes were stored at -80°C.

Samples were prepared similarly to TMT-labeled MHC-I peptides, however, instead of hipMHCs, 100 fmol of each of the 10 SIL-MHC complexes were added to each lysate. Unlabeled, lyophilized peptides eluted from the molecular weight cutoff filter were stored at -80°C until analysis. Peptides were resuspended in 5.25 μL 3% acetonitrile in 0.1% formic acid and 5 μL was injected for LC-MS/MS analysis.

### Sample preparation for unlabeled immunopeptidome analysis of GBM patient tumors

Tumor tissue was resuspended and homogenized in 750 μL to 1 mL of lysis buffer (20 mM Tris-HCl (pH 8.0), 150 mM NaCl, 0.2 mM phenylmethylsulfonyl fluoride (PMSF; Sigma), 1% CHAPS (Thermo Fisher), and 1× HALT Protease/Phosphatase Inhibitor Cocktail (Thermo Fisher). Lysates were cleared by centrifugation at 5,000 × g for 5 minutes at 4 °C, and protein concentration was measured by BCA assay.

The IP and subsequent filtering of peptides were performed as described for TMT-labeled MHC-I peptides. Multiple recurrence tumors (obtained from UCLA) were resuspended in 20 μL of 5% acetonitrile in 0.1% formic acid, and 5 μL was injected for LC-MS/MS analysis. First recurrence tumors (obtained from Mayo Clinic) underwent peptide fractionation. Lyophilized peptides were resuspended in 12 μL of 5% acetonitrile in 10 mM TEAB (pH 8.0) and loaded onto custom-packed C18 columns (200 μm inner diameter fused silica packed with 20 cm of 10 μm C18 beads, fritted on both ends). After sample loading, columns were rinsed with at least 50 μL of 5% acetonitrile in 10 mM TEAB. Samples were fractionated into 10 parts using high-pH reversed-phase chromatography on an Agilent 1100 Series HPLC with 10 mM TEAB and acetonitrile buffers. Fractions were lyophilized and stored at -80 °C. Before LC-MS/MS, fractions were resuspended in 5.25 μL of 5% acetonitrile in 0.1% formic acid, and 5 μL was injected for analysis.

### Gene Set Enrichment Analysis

Gene Set Enrichment Analysis was performed using the GSEApy Python package for GSEA (v4.3.3, RRID:SCR_025803).^24^ Pre-ranked GSEA was conducted against the Molecular Signatures Database (MSigDB) Hallmark gene sets, and Kyoto Encyclopedia of Genes and Genomes (KEGG) pathways where noted, using 1,000 permutations (seed = 6), with a minimum gene set size of 5 and maximum of 1,000.

For MHC-I peptides, where multiple peptides can originate from the same source protein, each peptide was mapped to its corresponding protein. Rankings were calculated based on the product of the log_2_ fold change and -log_10_ of the Benjamini-Hochberg adjusted p-value, based on TMT reporter ion intensities comparing TMZ-to DMSO-treated cells at 72 hours. Only peptides with non-zero TMT intensities in at least two replicates per condition were included. When multiple peptides mapped to the same source protein, the highest-ranking value was used for enrichment analysis.

### Statistical analysis and data visualization

Statistical analysis and data visualization were performed using Python (v3.7.9, RRID:SCR_008394), with key packages including NumPy (v1.21.6, RRID:SCR_008633), SciPy (v1.6.0, RRID:SCR_008058), scikit-learn (v0.23.2, RRID:SCR_002577), Matplotlib (v3.3.4, RRID:SCR_008624), and Seaborn (v0.11.1, RRID:SCR_018132). All t-tests were unpaired, two-sided Student’s t-tests, assuming equal variances and normal distribution, chosen given the balanced design with identical sample processing and multiplexed acquisition. No formal power analysis was performed. Sample sizes were determined by the maximum number of biological replicates that could be accommodated within a single TMT 18-plex experiment (three replicates per condition across six conditions and three timepoints). Benjamini-Hochberg correction was applied to control the false discovery rate at 5%.

### Data availability

The mass spectrometry data generated by this study are available via the ProteomeXchange Consortium (PRIDE, RRID:SCR_003411) using dataset identifier PXD067651 (DOI:10.6019/PXD067651). Processed transcripts per million from RNA sequencing can be found in Supplementary Table S3 and processed TMT intensities of translatome, proteome, phosphoproteome, and immunopeptidome data from LC-MS/MS can be found in Supplementary Table S4.

## Results

### Proteogenomic profiling reveals rapid, systems-level activation of the DNA damage response following TMZ exposure in GBM cells

To investigate the early effects of TMZ treatment, we performed an integrated, systems-level analysis of human U87-MG GBM cells. These cells have a methylated MGMT promoter yet exhibit moderate resistance to TMZ, with reported IC_50_ values between 58 and 650 µM.^25,26^ In initial experiments, 100 µM TMZ induced minimal cell death (Supplementary Fig. S1) while inducing the characteristic GC to AT mutational spectra associated with TMZ exposure (Supplementary Fig. S2).^27,28^ Thus, triplicate biological replicates of U87-MG cells were treated with either 100 µM TMZ or 0.1% DMSO (vehicle control) for 24, 48, and 72 hours. Given the asynchronous nature of cell cycling, we anticipated that although the reported doubling time of this cell line is ∼39 hours,^29^ a subset of cells would be undergoing division at each early time point, allowing the effect of TMZ to be observable even after shorter exposures. Because transcriptional changes do not uniformly correlate with protein abundance, translational activity, immune peptide presentation, or cell signaling states,^17,18,30–35^ we complemented bulk RNA sequencing with multiple parallel mass spectrometry-based methods to analyze each sample. Specifically, we used tandem mass tag (TMT)-based mass spectrometry to measure active protein translation using an improved method for quantifying newly synthesized proteins (translatome),^18^ quantify protein expression (proteome), assess kinase activity (phosphoproteome), and directly identify peptides presented on MHC-I molecules (immunopeptidome) with a recently developed approach from our laboratory (Fig. 1A).^10,19^

**Figure 1.**
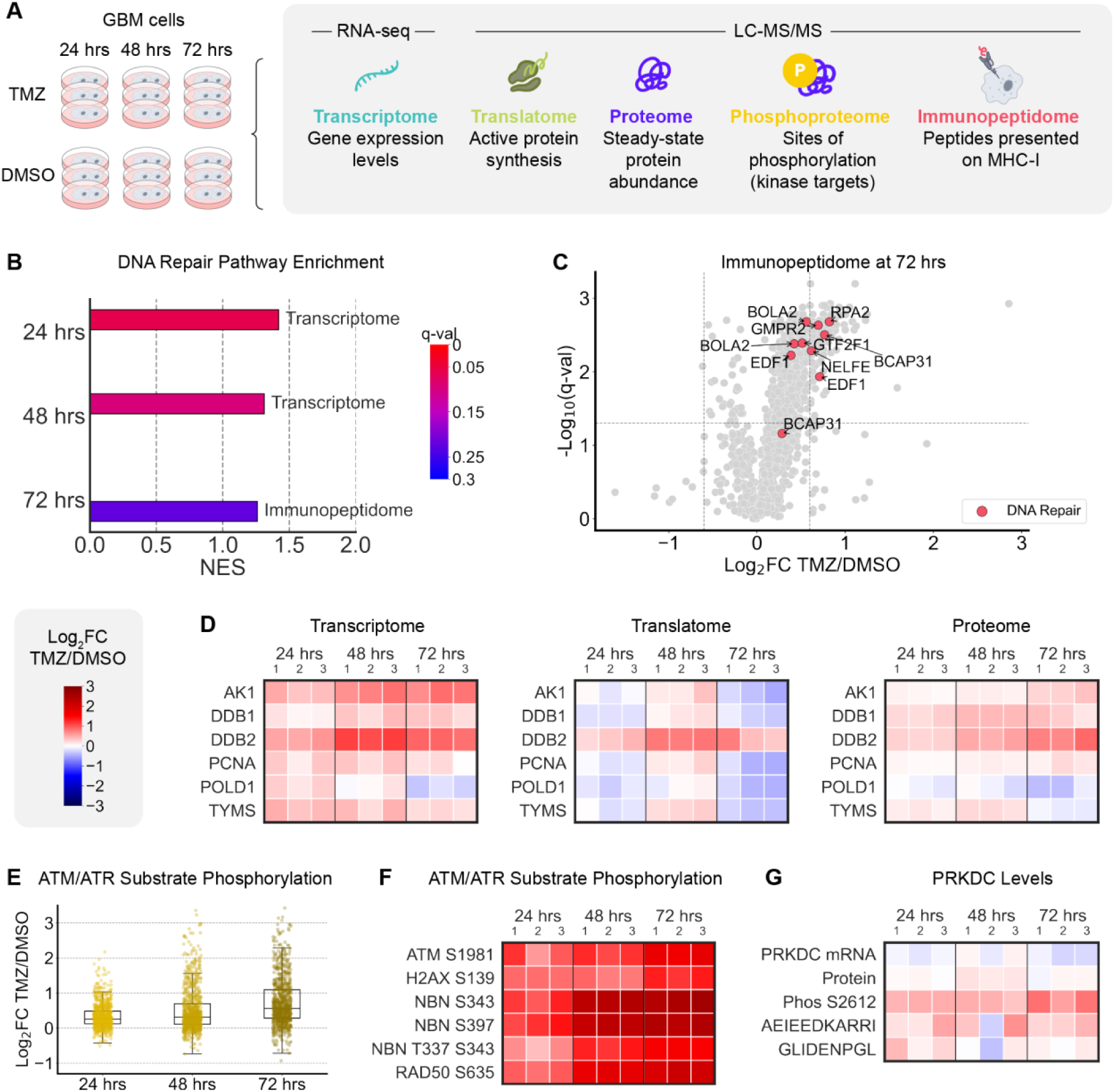
Integrated multi-omics analysis reveals early and coordinated activation of the DNA damage response following TMZ treatment in GBM cells. **A.** Overview of the multi-omics experimental workflow. U87-MG GBM cells were treated in triplicate with 100 µM TMZ or 0.01% DMSO for 24, 48, and 72 hours. Samples were analyzed using RNA sequencing (transcriptome) and TMT-labeled mass spectrometry to quantify newly synthesized proteins (translatome), protein abundance (proteome), phosphorylation of ATM/ATR and CDK kinase substrate motifs (phosphoproteome), and peptides presented on MHC-I molecules (immunopeptidome). **B.** Enrichment of the Hallmark DNA Repair gene set by pre-ranked GSEA with average log₂FC(TMZ/DMSO TPM or TMT signal) × –log₁₀(q-value) as the ranking metric. For the immunopeptidome, if multiple peptides mapped to the same source protein, the highest ranking was used. Normalized enrichment score (NES) | q-value: transcriptome at 24 hours = 1.421 | 0.061, 48 hours = 1.313 | 0.094; immunopeptidome at 72 hours = 1.261 | 0.224. **C.** Volcano plot of MHC-I peptides at 72 hours showing average log₂FC (TMZ/DMSO) vs. – log₁₀(q-value). Dotted lines indicate log₂FC = ±0.6 and q = 0.05. Hallmark DNA Repair pathway members are highlighted. **D.** Log_2_FC (TMZ/average DMSO TMT signal at each timepoint) of Hallmark DNA Repair gene set members consistently detected across transcriptome, translatome, and proteome datasets. **E.** Distribution of average log_2_FC (TMZ/DMSO TMT signal) phosphorylation of ATM/ATR substrate motifs (SQ or TQ) at each time point. Average log₂FC TMZ/DMSO: 24 hours = 0.341, 48 hours = 0.509, 72 hours = 0.738. **F.** Log₂FC (TMZ/average DMSO TMT signal at each timepoint) of canonical DNA damage markers at ATM/ATR substrate motifs. Average log_2_FC > 1 and q < 0.05 (unpaired two-sided t-test, FDR-adjusted) for all sites at all timepoints. **G.** Log_2_FC (TMZ/average DMSO TPM or TMT signal at each timepoint) of PRKDC (DNA-PKcs) transcript, protein, ABCDE cluster phosphorylation site, and two PRKDC-derived MHC-I peptides. The q-values were: PRKDC serine 2612 phosphorylation at 24 hours = 0.005, 48 hours = 0.022, 72 hours = 0.010; steady-state protein abundance at 48 hours = 0.007; PRKDC-derived MHC-I peptide AEIEEDKARRI at 72 hours = 0.043. Otherwise, no significant differences were observed, including in the transcriptome across all timepoints.

TMZ induces numerous potentially cytotoxic DNA lesions including methylation of guanine residues at the O^6^ position. The resulting O^6^-methylguanine lesion mis-pairs with thymine during replication, initiating futile cycles of mismatch repair (MMR), eventually leading to double-strand DNA breaks (DSBs).^1^ Despite the expectation that TMZ-induced DNA damage responses would require two cell cycles to become apparent, we detected activation of canonical DNA damage signaling pathways across multiple cellular regulatory levels within just 24 hours of exposure.

Pre-ranked gene set enrichment analysis (GSEA) of the RNA sequencing data using the ranking metric average log₂ fold-change (FC) of TMZ/average DMSO transcripts per million (TPM) at each timepoint × –log₁₀(Benjamini-Hochberg adjusted p-value from an unpaired, two-sided t-test of TMZ vs. DMSO) showed significant enrichment (q < 0.25) of the Hallmark DNA Repair gene set in the transcriptome at 24 and 48 hours (Fig. 1B). Intriguingly, this gene set was also significantly enriched in the immunopeptidome by 72 hours post-TMZ treatment, including an MHC-I peptide derived from RPA2, a replication protein A subunit involved in DNA replication and repair (Fig. 1C). These data indicate that the cellular response to TMZ may engage antigen presentation pathways to signal genomic instability to the immune system and that MHC-I peptides presented on the cell surface, to some extent, reflect internal cell processes.

Members of the Hallmark DNA Repair gene set detected across RNA sequencing, nascent protein translation, and global protein abundance datasets highlighted differential regulation of individual DNA repair-related genes (Fig. 1D). For example, AK1, DDB1, and PCNA exhibited increased gene expression upon TMZ treatment, coinciding with increased protein abundance, but not active translation, suggesting possible regulation via protein stability mechanisms. In contrast, thymidylate synthase (TYMS) exhibited an inverse relationship, with increased transcript levels but decreased protein abundance, suggesting translational suppression or enhanced degradation. Notably, DDB2, which recognizes and repairs UV-induced DNA lesions, showed consistent induction at the transcript, new protein, and protein levels following TMZ treatment as early as 24 hours, corroborating previous qRT-PCR observations in TMZ-treated U87-MG cells.^36^

Phosphoproteomic analysis revealed rapid DNA damage response activation within 24 hours of TMZ exposure, shown by increased ATM/ATR (Ataxia-Telangiectasia Mutated/ATM-and Rad3-related kinase) substrate motif phosphorylation (pSQ/pTQ) that progressively increased over three days (average log₂FC TMZ/DMSO TMT intensities: 24 hours = 0.341, 48 hours = 0.509, 72 hours = 0.738) (Fig. 1E). Among the proteins containing this phosphorylation motif were canonical DNA damage markers, including phosphorylated ATM at serine 1981, a well-characterized activating autophosphorylation site,^37^ and histone H2AX at serine 139, a marker of DNA double-stranded breaks.^38^ Additional ATM phosphorylation sites included nibrin (NBN) at serines 343 and 397, and RAD50 at serine 635, all of which play key roles in MRN complex-mediated DNA damage sensing and repair.^39–41^ These sites exhibited log_2_FC > 1 (TMZ/average DMSO TMT signal) at 24 hours, with increasing phosphorylation over time (Fig. 1F). By 72 hours, all markers had log_2_FC > 1 and FDR-adjusted p-values (q-value) < 0.01 (TMZ/DMSO, unpaired two-sided t-test, Benjamini-Hochberg correction).

DNA-PKcs (PRKDC), an essential component of non-homologous end joining (NHEJ), showed significantly increased phosphorylation at serine 2612 within 24 hours of TMZ treatment that was maintained through 72 hours (average log_2_FC | q-value: 24 hours = 0.466 | 0.005, 48 hours = 0.427 | 0.022, 72 hours = 0.740 | 0.010) (Fig. 1G). This site sits within the protein’s ABCDE cluster and is an autophosphorylation site critical for DNA-PKcs’s role in NHEJ; ATM is also known to phosphorylate this region.^42,43^ The observed increase in phosphorylation of this site suggests rapid activation of DNA-PK-mediated DNA repair. Increased phosphorylation was accompanied by moderate increases in PRKDC protein abundance (particularly at 48 hours) and one PRKDC-derived MHC-I peptide, despite no significant change in transcript levels, suggesting that post-translational modifications are key in regulating DNA-PKcs activity in response to TMZ.

Together, these results indicate that TMZ rapidly triggers a multilayered DNA damage response in GBM cells within 24 hours following TMZ exposure.

### TMZ induces systems-level activation of p53 signaling and cell cycle arrest

In the canonical DNA damage response pathway, DNA damage signaling leads to p53 activation, which can drive apoptosis or cell cycle arrest to enable DNA repair. Accordingly, alongside the DNA damage response, TMZ treatment rapidly activated the p53 signaling pathway, with significant enrichment (q < 0.25) of the Hallmark p53 Pathway detected by pre-ranked GSEA in the transcriptome at all timepoints, in the proteome at 72 hours, and in the immunopeptidome at 24 and 72 hours (Fig. 2A). MHC-I peptides derived from p53 Pathway members with increased presentation at 72 hours post-TMZ treatment included those from GADD45A and PLK2, both involved in cell cycle regulation, and HMOX1, a stress response protein induced under oxidative and genotoxic stress conditions (Fig. 2B). These data suggest that internal p53 signaling is communicated to the immune system through altered immunopeptide presentation, potentially enabling immune recognition and elimination of DNA-damaged cells.

**Figure 2.**
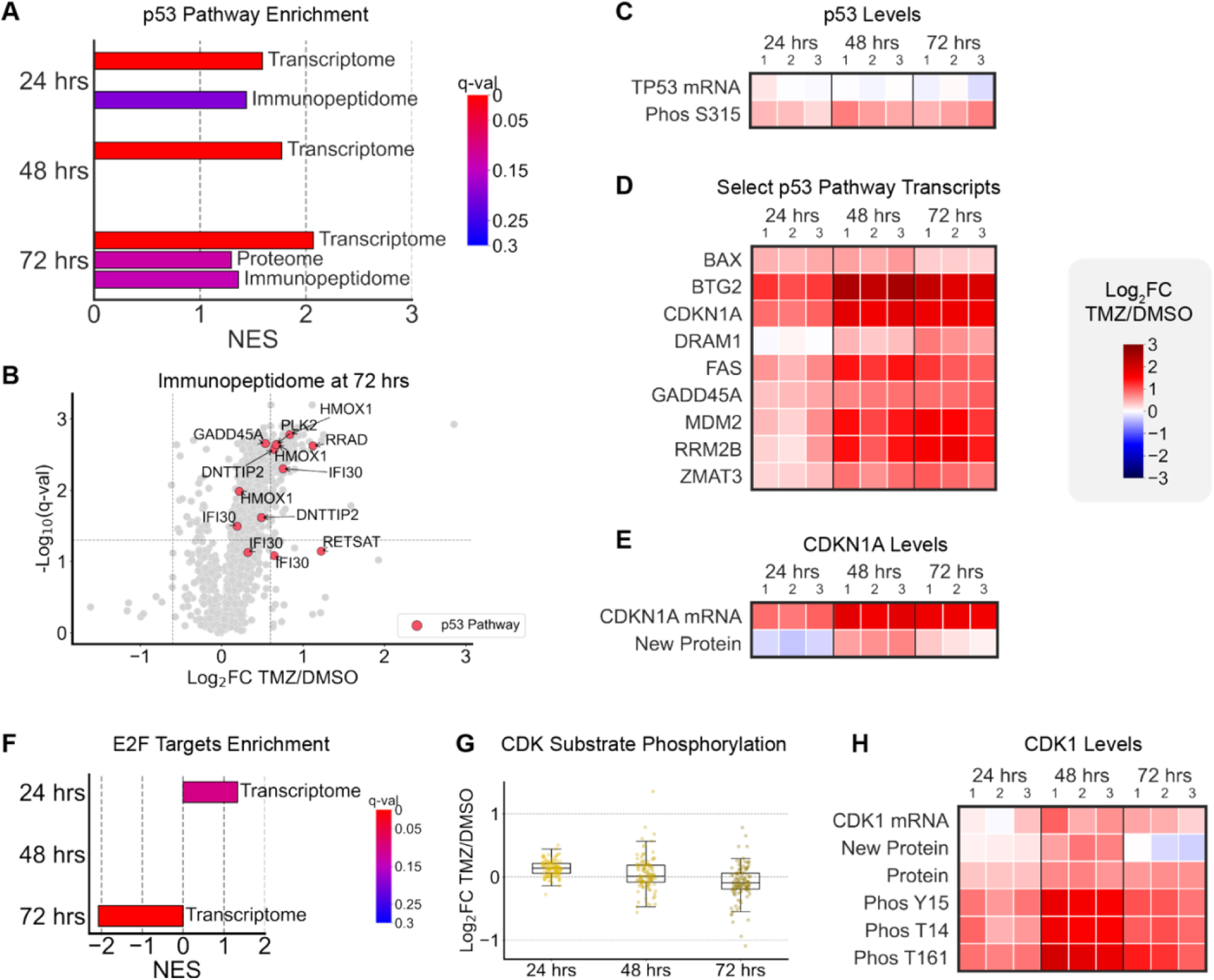
TMZ treatment rapidly activates the p53 pathway and cell cycle arrest. **A.** Enrichment of the Hallmark p53 Pathway gene set by pre-ranked GSEA with average log₂FC(TMZ/DMSO TPM or TMT signal) × –log₁₀(q-value) as the ranking metric. For the immunopeptidome, if multiple peptides mapped to the same source protein, the highest ranking was used. Normalized enrichment score (NES) | q-value: transcriptome at 24 hours = 1.590 | 0.001, 48 hours = 1.772 | 0.000, 72 hours = 2.069 | 0.000; proteome at 72 hours = 1.294 | 0.128; immunopeptidome at 24 hours = 1.437 | 0.199, 72 hours = 1.363 | 0.141. **B.** Volcano plot of MHC-I peptides at 72 hours showing average log₂FC (TMZ/DMSO) vs. – log₁₀(q-value). Dotted lines indicate log₂FC = ±0.6 and q = 0.05. Hallmark p53 Pathway members are highlighted. **C.** Log_2_FC (TMZ/average DMSO TPM or TMT signal at each timepoint) of TP53 transcript and serine 315 phosphorylation site. For S315, the q-values were: 24 hours = 0.316; 48 hours = 0.050; 72 hours = 0.077. No significant differences were observed in TP53 transcript levels. **D.** Log₂FC (TMZ/average DMSO TPM at each timepoint) of well-characterized p53 pathway genes ranking in the top 10% most differentially expressed at 48 and 72 hours (except BAX: top 10% at 48 hours, top 20% at 72 hours) based on log₂FC × –log₁₀(q-value). **E.** Log₂FC (TMZ/average DMSO TPM or TMT signal at each timepoint) of CDKN1A transcript and newly synthesized protein. For the transcript, q-values were: 24 hours = 0.348; 48 hours = 0.035; 72 hours = 0.036. No significant differences were observed in CDKN1A newly synthesized protein levels. **F.** Enrichment of the Hallmark E2F Targets gene set by pre-ranked GSEA with average log₂FC(TMZ/DMSO TPMs) × –log₁₀(q-value) as the ranking metric. Normalized enrichment score (NES) | q-value: transcriptome at 24 hours = 1.335 | 0.105, 72 hours = -2.073 | 0.000. **G.** Distribution of average log_2_FC (TMZ/DMSO TMT signal) phosphorylation of CDK substrate motifs (SPXK, TPXK, SPXR, or TPXR) at each time point. Average log₂FC TMZ/DMSO: 24 hours = 0.135, 48 hours = 0.051, 72 hours = -0.087. **H.** Log_2_FC (TMZ/average DMSO TPMs or TMT signal at each timepoint) of CDK1 transcript, newly synthesized protein, steady-state protein, and well-characterized activating and deactivating phosphorylation sites. The q-values were: protein at 24 hours = 0.011, 48 hours = 0.006, 72 hours = 0.005; tyrosine 15 phosphorylation at 24 hours = 0.107, 48 hours = 0.009, 72 hours = 0.016; threonine 14 phosphorylation at 24 hours = 0.264, 48 hours = 0.009, 72 hours = 0.017; threonine 161 phosphorylation at 24 hours = 0.028, 48 hours = 0.010, 72 hours = 0.037. No significant differences were observed in transcript or newly synthesized protein levels.

Additionally, p53 activity was detectable by increased phosphorylation of TP53 at serine 315, a residue within the DNA-binding domain implicated in cell cycle control that allows cells to arrest and repair damage before mitosis.^44^ Unlike the TP53 transcript, which showed no significant TMZ-associated changes, the phosphorylation site significantly increased with TMZ at 48 hours and nearly significantly at 72 hours (average log_2_FC | q-value: 48 hours = 0.618 | 0.050, 72 hours = 0.581 | 0.077) (Fig. 2C).

Several Hallmark p53 Pathway genes, including canonical p53 targets involved in apoptosis, DNA repair, and growth arrest, ranked in the top 10% most differentially expressed genes (based on log₂FC × –log₁₀(q-value)) at 48 and 72 hours (Fig. 2D). These genes included BAX (top 10% at 48 hours, top 20% at 72 hours), BTG2, DRAM1, FAS, GADD45A, RRM2B, and ZMAT3. Notably, MDM2, a transcriptional target and negative feedback regulator of p53, was also strongly upregulated, along with CDKN1A, a direct downstream effector of MDM2. CDKN1A, which encodes the cyclin-dependent kinase inhibitor p21 and is a key mediator of p53-driven G1/S arrest, showed one of the largest increases in gene expression in response to TMZ at each time point (average log_2_FC | q-value: 24 hours = 0.856 | 0.348, 48 hours = 1.770 | 0.035, 72 hours = 1.677 | 0.036) (Fig. 2E). However, its translation only modestly, but not significantly, increased with TMZ over time (average log_2_FC | q-value: 24 hours = -0.265 | 0.056, 48 hours = 0.634 | 0.997, 72 hours = 0.195 | 0.215) (Fig. 2E). This discrepancy may reflect transcriptional priming without full engagement of translational machinery at these time points, or it may result from dynamic range compression inherent to TMT-based mass spectrometry, which can obscure changes in protein abundance.^45^

Further evidence of TMZ-mediated cell cycle regulation emerged from the enrichment of the Hallmark E2F Targets gene set in the transcriptome (Fig. 2F). This pathway was significantly (q = 0.105) positively enriched at 24 hours, likely reflecting early upregulation of DNA repair genes (many of which are E2F targets). It then became significantly (q = 0.000) negatively enriched by 72 hours, consistent with a shift toward cell cycle arrest. This shift was further corroborated by phosphoproteomic evidence of reduced cyclin-dependent kinase (CDK) activity over time with TMZ, as indicated by decreased overall phosphorylation of known CDK substrate motifs (pS/T-PXK and pS/T-PXR) (average log₂FC TMZ/DMSO TMT intensities: 24 hours = 0.135, 48 hours = 0.051, 72 hours = -0.087) (Fig. 2G).

Additionally, while CDK1 mRNA and nascent protein synthesis did not significantly change, TMZ significantly increased total CDK1 protein abundance (average log_2_FC | q-value: 24 hours = 0.350 | 0.011, 48 hours = 0.687 | 0.006, 72 hours = 0.627 | 0.005) and phosphorylation of key sites (Fig. 2H). Phosphorylation of CDK1 at T14 and Y15, two established inhibitory sites that suppress CDK1 activity, and T161, an activating site required for CDK1 priming, all significantly increased with TMZ over time (average log₂FC > 0.97 and q < 0.05 for all sites at 48 and 72 hours) (Fig. 2H). Phosphopeptides corresponding to these sites were identified as IGEGTpYGVVYK (Y15) and IGEGpTYGVVYK (T14), both of which are conserved across CDK1, CDK2, and CDK3, and thus cannot be unambiguously assigned. However, given the biological context and the critical role of CDK1 in G2/M checkpoint control, these signals likely reflect CDK1 regulation. Phosphorylation at T161 was detected via the CDK1-specific phosphopeptide VYpTHEVVTLWYR and is known to co-occur with T14, suggesting a checkpoint-enforced mechanism that permits priming without activation.^46^ Thus, while CDK1 appears biochemically prepared for mitotic entry, TMZ-induced DNA damage could override its activation, resulting in suppression of CDK activity. However, it is also possible that these changes reflect an accumulation of cells in G2/M rather than per cell upregulation of CDK1.

### TMZ induces a rapid and multilayered cell stress response

In addition to eliciting a DNA damage response, TMZ is known to trigger a spectrum of cellular stress pathways in GBM, many of which are associated with therapeutic resistance.^47–49^ In our study, evidence of oxidative and metabolic stress were evident across multiple cellular regulatory layers within the first few days of treatment.

In the transcriptome, several Hallmark stress-associated gene sets, including Xenobiotic Metabolism, TNF-α signaling via NF-κB, IL-6/JAK/STAT3 signaling, Inflammatory Response, and Oxidative Phosphorylation were significantly enriched (q < 0.25), particularly at the 72-hour time point (Fig. 3A). The top 10% most differentially expressed genes at 72 hours included members of these gene sets that are well-established markers of acute cellular stress: ATF3, DDIT3, IER3, PTGS2 (COX-2), KYNU, NAMPT, and RCAN1 (Fig. 3B). Interestingly, other classic stress markers such as HMOX1, HSPA1A, and HSPA1B showed no notable transcriptional upregulation following TMZ treatment (Fig. 3B).

**Figure 3.**
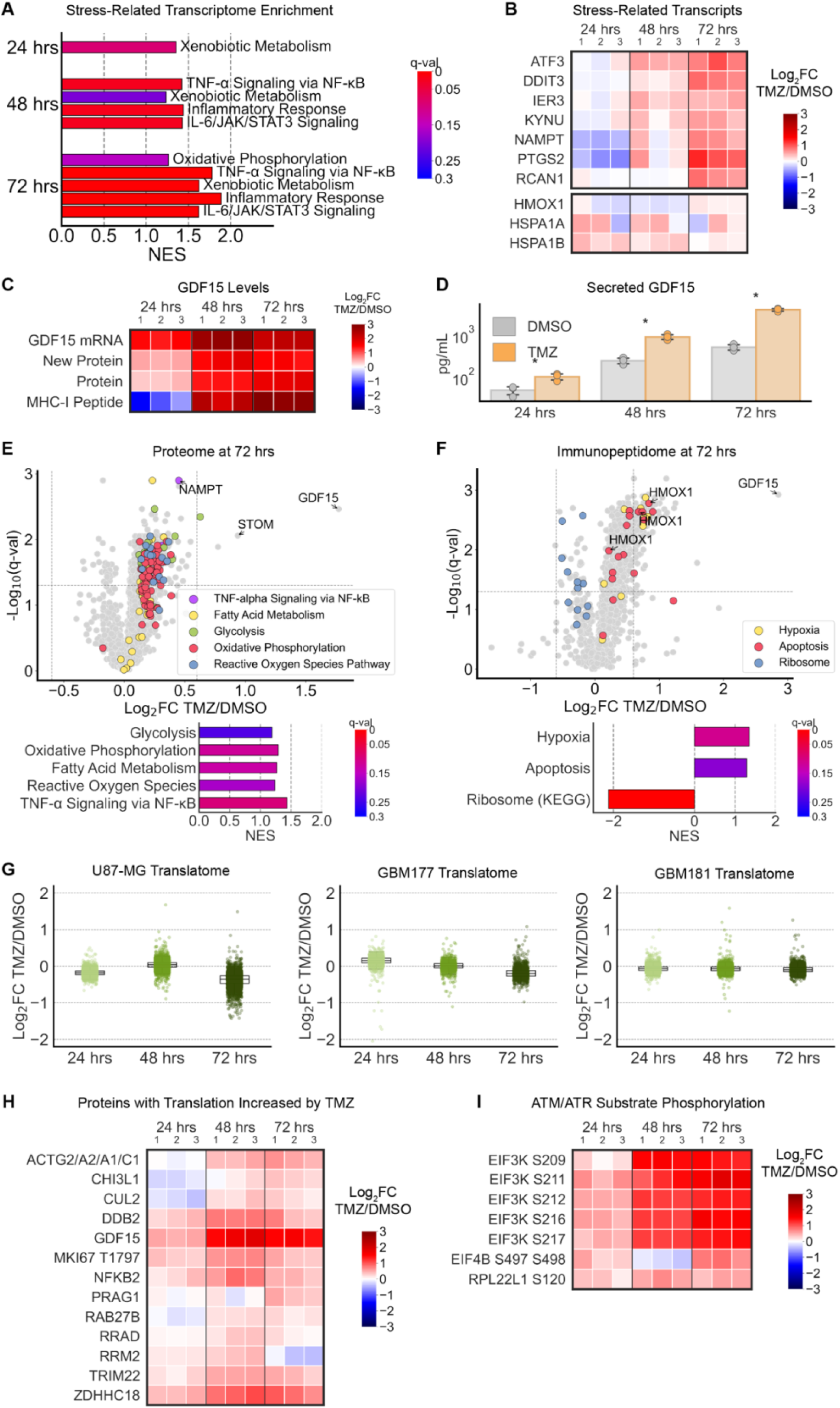
TMZ rapidly induces stress responses across multiple regulatory layers, including suppression of translation. **A.** Enrichment of stress-related Hallmark gene sets in the transcriptome by pre-ranked GSEA with average log₂FC(TMZ/DMSO TPM) × –log₁₀(q-value) as the ranking metric. Normalized enrichment score (NES) | q-value: Xenobiotic Metabolism at 24 hours = 1.354 | 0.090, 48 hours = 1.235 | 0.204, 72 hours = 1.626 | 0.003; TNF-α Signaling via NF-κB at 48 hours = 1.422 | 0.024, 72 hours = 1.781 | 0.000; IL-6/JAK/STAT3 Signaling at 48 hours = 1.428 | 0.024, 72 hours = 1.622 | 0.003; Oxidative Phosphorylation at 72 hours = 1.264 | 0.155; Inflammatory Response at 72 hours = 1.888 | 0.000. **B.** Log₂FC (TMZ/average DMSO TPM at each timepoint) of stress-related genes (from pathways in A) ranking in the top 10% most differentially expressed at 72 hours based on log₂FC × – log₁₀(q-value). Shown alongside HMOX1, HSPA1A, and HSPA1B, stress-related transcripts that did not meet the 10% cutoff. **C.** Log_2_FC (TMZ/average DMSO TPM or TMT signal at each timepoint) of GDF15 transcript, newly synthesized protein, steady-state protein, and ALPEGLPEA MHC-I peptide. The q-values were: transcript at 24 hours = 0.348, 48 hours = 0.028, 72 hours = 0.036; newly-synthesized protein at 24 hours = 0.046, 48 hours = 0.717, 72 hours = 0.030; steady-state protein at 24 hours = 0.036, 48 hours = 0.000, 72 hours = 0.003; MHC-I peptide at 24 hours = 0.999, 48 hours = 0.259, 72 hours = 0.001. **D.** Concentration (pg/mL) of secreted GDF15 in cell culture media measured by ELISA. Asterisks indicate q < 0.05 by unpaired two-sided t-test. Average concentration DMSO vs. TMZ: 24 hours = 49.90 vs. 105.1 pg/mL, q = 0.018; 48 hours = 251.5 vs. 916.0 pg/mL, q = 0.002; 72 hours = 524.9 vs. 4,015 pg/mL, q = 0.000. **E.** Volcano plot of proteins at 72 hours showing average log₂FC (TMZ/DMSO) vs. –log₁₀(q-value). Dotted lines indicate log₂FC = ±0.6 and q-value = 0.05. Stress-related Hallmark gene sets are highlighted and enrichment shown in the barplot (pre-ranked GSEA with average log₂FC(TMZ/DMSO TPM) × –log₁₀(q-value) as the ranking metric). Normalized enrichment score (NES) | q-value at 72 hours: Glycolysis = 1.190 | 0.237, Oxidative Phosphorylation = 1.293 | 0.119, Fatty Acid Metabolism = 1.265 | 0.130, Reactive Oxygen Species Pathway = 1.236 | 0.168, TNF-α Signaling via NF-κB = 1.434 | 0.094. **F.** Volcano plot of MHC-I peptides at 72 hours showing average log₂FC (TMZ/DMSO) vs. – log₁₀(q-value). Dotted lines indicate log₂FC = ±0.6 and q = 0.05. Stress-related Hallmark gene sets are highlighted and enrichment shown in the barplot (pre-ranked GSEA with average log₂FC(TMZ/DMSO TPM) × –log₁₀(q-value) as the ranking metric; if multiple peptides mapped to the same source protein, the highest ranking was used). Normalized enrichment score (NES) | q-value at 72 hours: Hypoxia = 1.354 | 0.110, Apoptosis = 1.291 | 0.192, Ribosome (KEGG) = -2.120 | 0.000. **G.** Distribution of average log_2_FC (TMZ/DMSO TMT signal) newly synthesized proteins at each time point in U87-MG and partially HLA-matched primary glioma lines. Average log_2_FC of all proteins at 72 hours: U87-MG = -0.374, GBM177 = -0.184, GBM181 = -0.087. **H.** Log₂FC (TMZ/average DMSO TMT signal) of newly synthesized proteins with average log2FC > 0 and q < 0.1 at any timepoint following TMZ treatment. One peptide was derived from a conserved region of muscle actins (ACTG2, ACTA2, ACTA1, ACTC1), preventing assignment to a specific isoform. **I.** Log₂FC (TMZ/average DMSO TMT signal at each timepoint) of translation-related proteins at ATM/ATR substrate motifs. Average log_2_FC > 0.4 and q-value < 0.02 (unpaired two-sided t-test, FDR-adjusted) for all sites at 72 hours.

Among the most striking markers of TMZ-induced stress was GDF15, which was robustly induced across all measured levels of regulation (Fig. 3C). GDF15 is a member of the TGF-β cytokine superfamily and a known p53 transcriptional target that is upregulated in response to acute stress, and also contributes to tumor progression, adaptation, and immune evasion in gliomas.^50,51^ Additionally, it has been characterized as a secreted marker of the integrated stress response in the central nervous system.^52^ By 72 hours post-treatment, the GDF15 transcript, nascent protein, protein, and corresponding MHC-I peptide were all significantly elevated (log_2_FC > 1.48 and q < 0.05), with the nascent protein and protein exhibiting q < 0.05 as early as 24 hours. ELISA confirmed significantly elevated GDF15 secretion with increasing levels over time (average concentration DMSO vs. TMZ: 24 hours = 49.90 vs. 105.1 pg/mL, q = 0.018; 48 hours = 251.5 vs. 916.0 pg/mL, q = 0.002; 72 hours = 524.9 vs. 4015 pg/mL, q = 0.0002) (Fig. 3D).

Consistent with the energy demands of DNA repair and survival under stress, TMZ treatment led to enrichment of proteomic pathways associated with cellular metabolism. Hallmark gene sets for Glycolysis, Oxidative Phosphorylation, Fatty Acid Metabolism, and Reactive Oxygen Species (ROS) Pathway were all significantly enriched (q < 0.25) in the proteome at 72 hours (Fig. 3E). This metabolic shift may be driven by upstream activation of p53 and ATM/ATR kinases, which can induce ROS production as a signaling mechanism. TNF-α signaling via NF-κB, an inflammatory pathway downstream of DNA damage and oxidative stress, was also enriched in the proteome, further connecting stress signaling to metabolic reprogramming.

The immunopeptidome also reflected signatures of cellular stress (Fig. 3F). Peptides derived from proteins in the Hypoxia and Glycolysis gene sets were significantly enriched (q < 0.25) at 24 hours post-TMZ treatment. By 72 hours, Hypoxia and Apoptosis pathways were significantly enriched (Glycolysis q = 0.281) (Fig. 3F). Because cells were grown under normoxic conditions, the Hypoxia pathway likely reflects stress signaling rather than low oxygen. These data further suggest that the intracellular consequences of DNA damage and oxidative stress are reflected on the cell surface through MHC-I peptide presentation. In contrast, the KEGG Ribosome gene set was negatively enriched in the immunopeptidome, with several ribosome-associated peptides showing decreased MHC-I presentation at 72 hours, suggesting stress-induced translational suppression or altered protein turnover in response to TMZ (Fig. 3F).

### Cell stress in response to TMZ is associated with decreased translation

While transcriptomic and proteomic analyses revealed changes associated with DNA damage, p53 activation, and cellular stress, these methods alone may miss the rapid and dynamic shifts in translation that occur in response to TMZ. Translational control plays a distinct and critical role in shaping the adaptive response to genotoxic stress, and it has been previously linked to DNA damage, tumorigenesis, and antigen processing.^33,53–56^ Importantly, translation is a distinct process that does not necessarily correlate with transcription or steady-state protein levels,^18^ making the translatome a source of additional insight into the early adaptive responses to TMZ and its impact on immune recognition. Additionally, p53 is known to suppress translation through mTOR under stress conditions.^57^

To directly measure changes in nascent protein synthesis following TMZ exposure, we applied an improved version of the multiplex isobaric tagging/non-canonical amino acid tagging (MITNCAT) method,^18^ which labels actively translated proteins during a one-hour window with amino acid analogs. We enhanced the quantitation accuracy of this method by incorporating synthetic heavy isotope labeled peptide standards. This refined MITNCAT approach revealed that, while translation changes were subtle during the first two days of TMZ treatment, by 72 hours, global translation of approximately 3,000 newly synthesized proteins was reduced in U87-MG cells relative to DMSO controls (average log_2_FC of all proteins at 72 hours = -0.374) (Fig. 3G). The global reduction in protein synthesis coincided with the reduced presentation of ribosomal protein-derived peptides in the immunopeptidome (Fig. 3F). Post-TMZ decreased translation was also observed in two primary MGMT methylated glioma lines, GBM177 and GBM181, partially HLA-matched to MGMT methylated U87-MG (U87-MG alleles HLA-A*02:01, HLA-B*44:02, HLA-C*05:01; GBM177 alleles HLA-A*02:01/03:01, HLA-B*07:02/51:01, HLA-C*07:02/14:02; GBM181 alleles HLA-A*02:01/26:01, HLA-B*44:02/51:01, HLA-C*02:02/05:01), though the magnitude of suppression was less pronounced (average log_2_FC of all proteins at 72 hours: GBM177 = -0.184, GBM181 = -0.087), likely reflecting their slower doubling times (Fig. 3G).

Of the ∼3,000 newly translated proteins detected, only 13 showed appreciably increased translation (average log2FC > 0 and q < 0.1) at any timepoint following TMZ treatment (Fig. 3H). All 13 proteins have been implicated in cancer progression, survival, and/or treatment resistance. Some have established roles in ubiquitination and protein degradation, such as CUL2 (E3 ubiquitin ligase component), DDB2 (NA damage-binding protein that complexes with a ubiquitin ligase), and TRIM22 (RING-type E3 ubiquitin ligase). The most significantly upregulated proteins were GDF15 and ZDHHC18 at 72 hours (average log_2_FC | q-value: GDF15 = 1.488 | 0.030, ZDHHC18 = 0.842 | 0.031). GDF15, beyond its cellular stress response role, is upregulated by radiation in GBM and contributes to radioresistance in glioma stem cells and breast cancer.^58–60^ ZDHHC18, a palmitoyl transferase, negatively regulates cGAS-mediated innate immune responses and has been implicated in GBM tumorigenesis.^61,62^ Its increased translation may enable immune evasion despite ongoing TMZ-induced DNA damage. Notably, we detected increased translation of a phosphorylated MKI75 (Ki-67) peptide at 24 hours (average log_2_FC | q-value: 0.316 | 0.047), while unphosphorylated Ki-67 new protein and mRNA levels remained unchanged or decreased (q-values > 0.1 at all time points except at 72 hours, when newly translated Ki-67 decreased with average log_2_FC | q-value: -0.468 | 0.035). Although this phosphorylation site is not well characterized, its selective induction suggests a potential non-canonical role for Ki-67 during TMZ-induced stress. Beyond its function as a proliferation marker, Ki-67 protects DNA replication forks and maintains genome stability,^63^ potentially facilitating adaptive chromatin or replication responses under chemotherapeutic stress. The selective upregulation of this small set of proteins amidst broad translational suppression suggests a stress-adaptive reprogramming of protein synthesis that prioritizes cell survival and early resistance.

Phosphoproteomic analysis revealed significantly increased ATM/ATR-mediated phosphorylation of translation elongation factors (EIF3K, EIF4B) and ribosomal protein RPL22L1 in TMZ-treated cells (average log2FC > 0.4 and q < 0.02 at 72 hours for all peptides shown in Fig. 3I). In contrast, phosphorylation by non-ATM/ATR kinases on these proteins did not significantly increase (except site EIF4B serine 597) (Supplementary Fig. S3), suggesting that DNA damage specifically modifies components of the translation machinery. While the functional roles of these specific phosphosites are largely uncharacterized, analogous modifications, such as phosphorylation of eIF2α, are known to suppress translation initiation.^64,65^

### TMZ activates interferon gamma response and antigen presentation machinery

Beyond initiating DNA repair, p53-related, and cell stress responses, TMZ treatment also activated components of the interferon gamma (IFN-γ) signaling pathway and enhanced antigen presentation. DNA damage is known to trigger type I interferon and cytokine genes via activation of the cGAS-STING pathway, which can stimulate immune activation.^4,5,66,67^ The Hallmark Interferon Gamma Pathway gene set was significantly enriched (q < 0.25) in both the transcriptome (at 48 and 72 hours) and proteome (at 72 hours) following TMZ treatment (Fig. 4A), including upregulation of numerous components involved in antigen processing and presentation. Among the top 10% most differentially expressed genes at 72 hours were multiple key MHC-I complex components, such as B2M and multiple HLA loci, and peptide transporter TAP1 (Fig. 4B). Genes FDXR (p53-responsive gene involved in IFN-γ regulation), IL6 (pro-inflammatory cytokine), and SESN1 (53 and oxidative stress target implicated in cytokine signaling), showed significantly increased expression (log_2_FC > 1 and q < 0.05) by 72 hours post-TMZ. Protein-level increases in IFN-γ and antigen presentation machinery were evident by 72 hours (Fig. 4C). Several components of the MHC-I assembly and presentation pathway were among the top 10% most differentially abundant proteins at 72 hours, including CANX (calnexin) and CALR (calreticulin), which facilitate MHC-I folding and peptide loading; proteasomal subunits PSME1 and PSME2; and UBE2L6, an E2 ubiquitin-conjugating enzyme that supports protein degradation for antigen processing. At the transcript level, HLA-A and CANX ranked in the top 13% and 17% of differentially expressed genes, respectively.

**Figure 4.**
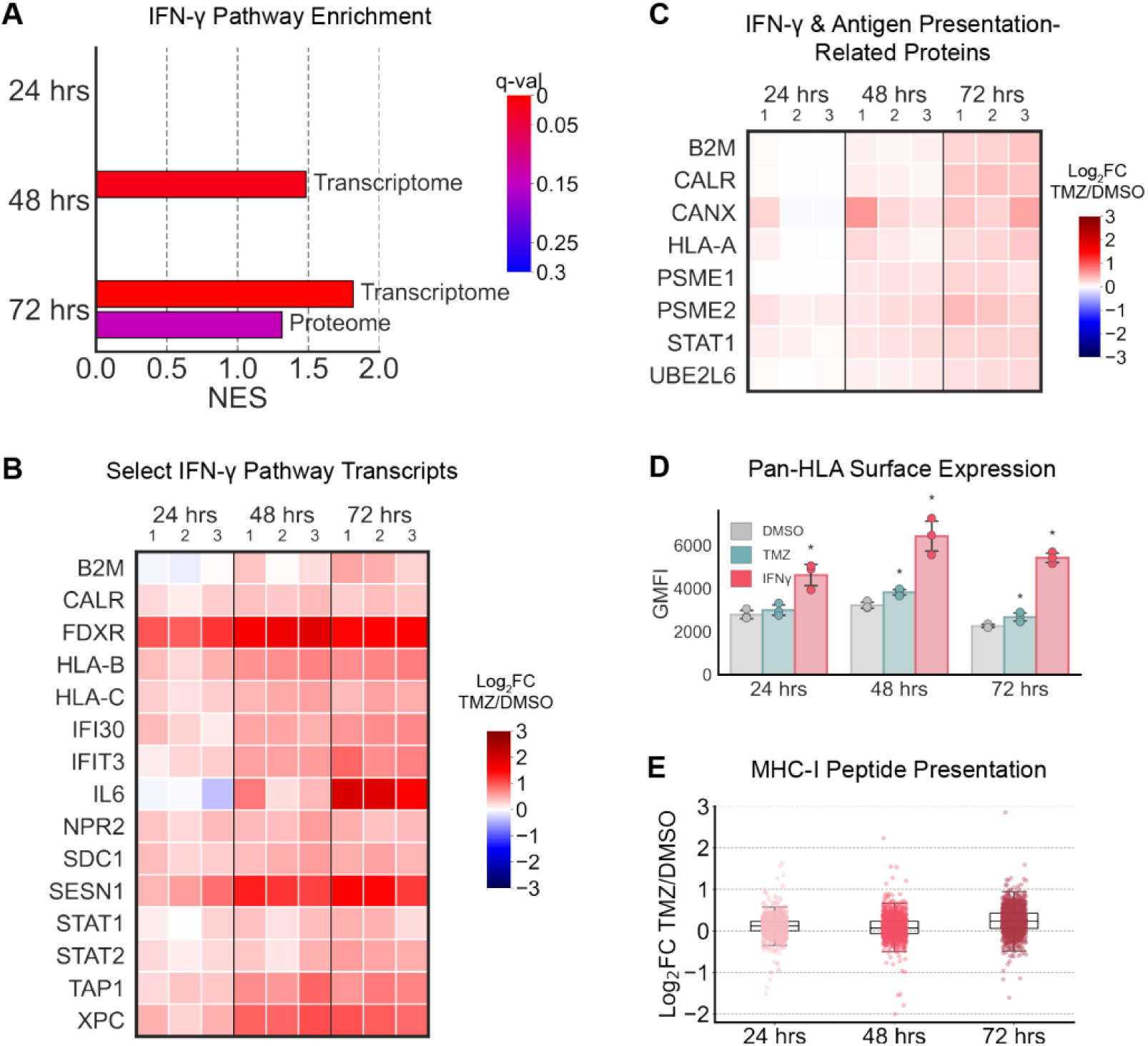
TMZ activates interferon gamma responses and antigen presentation in GBM cells. **A.** Enrichment of the Hallmark Interferon Gamma Pathway gene set by pre-ranked GSEA with average log₂FC(TMZ/DMSO TPM or TMT signal) × –log₁₀(q-value) as the ranking metric. Normalized enrichment score (NES) | q-value: transcriptome at 48 hours = 1.484 | 0.018, 72 hours = 1.817 | 0.000; proteome at 72 hours = 1.314 | 0.142. **B.** Log₂FC (TMZ/average DMSO TPM at each timepoint) of well-characterized Interferon Gamma Pathway genes ranking in the top 10% most differentially expressed at 72 hours based on log₂FC × –log₁₀(q-value). **C.** Log₂FC (TMZ/average DMSO TMT signal at each timepoint) of well-characterized IFN-γ and antigen presentation-related proteins from the proteome ranking in the top 10% most differentially abundant at 72 hours based on log₂FC × –log₁₀(q-value). **D.** Geometric mean fluorescence intensity (GMFI) of surface pan-HLA molecules measured by flow cytometry. Asterisks indicate q < 0.05 by unpaired two-sided t-test compared to DMSO. Average GMFI values DMSO vs. TMZ vs. IFN-γ: 24 hours = 2,788 vs. 3,001 vs. 4,627; 48 hours = 3,225 vs. 3,832 vs. 6,430; 72 hours = 2,269 vs. 2,679 vs. 5,424. **E.** Distribution of average log_2_FC (TMZ/DMSO TMT signal) MHC-I peptides at each time point. Average log₂FC TMZ/DMSO: 24 hours = 0.111, 48 hours = 0.086, 72 hours = 0.240.

Flow cytometry using a pan-HLA antibody confirmed a progressive increase in surface MHC-I levels over time, with significant differences between TMZ and DMSO treatments emerging by 48 hours and increasing further at 72 hours (Fig. 4D, Supplementary Fig. S4).

To assess the functional consequences of these changes, we profiled the repertoire of MHC-I peptides at each time point. Although the overall distribution of peptide-level log_2_FC subtly shifted over time with TMZ (average log₂FC TMZ/DMSO TMT intensities: 24 hours = 0.111, 48 hours = 0.086, 72 hours = 0.240) (Fig. 4E), specific subsets of peptides were differentially presented. As early as 24 hours, pre-ranked GSEA of MHC-I peptides revealed significant (q < 0.25) enrichment of the p53 Pathway, Hypoxia, Epithelial to Mesenchymal Transition (EMT), and Glycolysis Hallmark gene sets (Supplementary Table S1). By 72 hours, p53 Pathway and Hypoxia remained enriched while Apoptosis and DNA Repair emerged, indicating that antigen presentation dynamically evolves to reflect the shifting landscape of intracellular stress and DNA damage (Supplementary Table S1).

### TMZ increases presentation of treatment-associated peptide antigens (TAPAs)

By 72 hours post-treatment, TMZ induced a shift in the MHC-I immunopeptidome, with a subset of peptides showing robust and consistent increases in presentation over time. Applying cutoffs of log_2_FC > 0.75 and q < 0.05 at 72 hours nominated 51 candidate treatment-associated peptide antigens (TAPAs). From these, we selected 10 representative peptides for experimental validation based on the biological relevance of their source proteins to TMZ-induced cellular processes (Fig. 5A). These candidate TAPAs were derived from proteins involved in: DNA damage and repair (PARP1, TP53BP1), stress-responsive cytokine signaling (GDF15, CSF3), immune-related processing and presentation (IFI30), ROS regulation and metabolic adaptation (RETSAT, AMPD2), matrix remodeling (HAS1), mitochondrial transport (SLC25A35), and proteasomal degradation (UBAP2L). Two peptides were included despite slightly missing the thresholds: a RETSAT-derived peptide (q = 0.071) was retained given RETSAT’s role in oxidative stress regulation and ROS generation,^68^ and a phosphorylated AMPD2-derived peptide (log_2_FC = 0.454) was included based on the potential for phosphopeptides to serve as neoantigens. These post-translationally modified peptides may arise from dysregulated signaling in tumor cells and therefore could confer tumor specificity; moreover, phosphorylation can enhance MHC-I binding affinity to create unique antigenic forms capable of eliciting T cell responses.^69–71^ In addition to the phosphorylated AMPD2-derived peptide, we also included a phosphorylated TP53BP1-derived peptide (log_2_FC = 0.776, q = 0.002).

**Figure 5.**
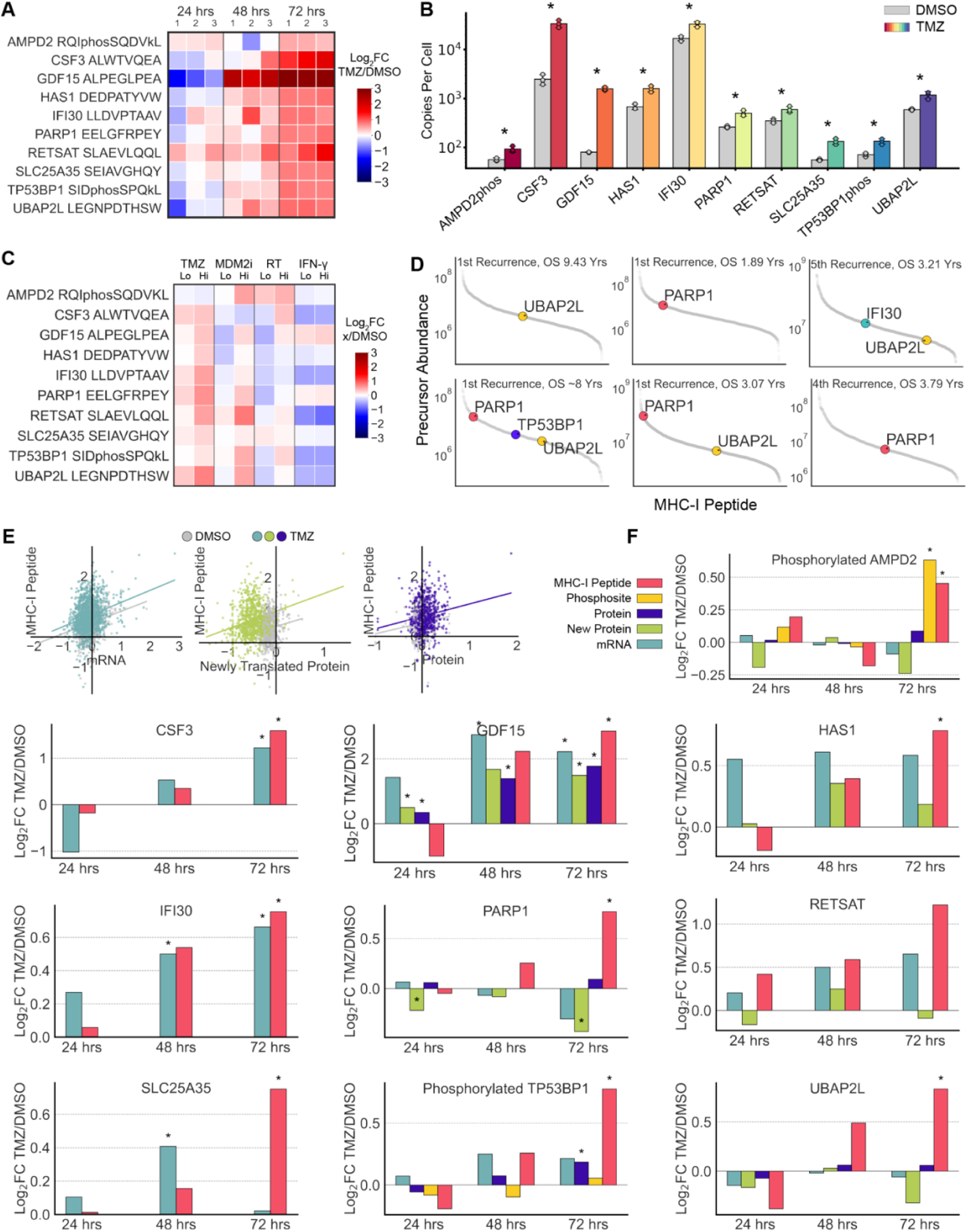
TMZ increases presentation of novel treatment-associated peptide antigens. **A.** Log₂FC (TMZ/average DMSO TMT signal at each timepoint) of candidate TMZ treatment-associated peptide antigens (TAPAs). At 72 hours, log_2_FC | q-value: phosphorylated AMPD2 = 0.454 | 0.011; CSF3 = 1.590 | 0.016; GDF15 = 2.854 | 0.001; HAS1 = 0.786 | 0.001; IFI30 = 0.755 | 0.005; PARP1 = 0.769 | 0.002; RETSAT = 1.222 | 0.071; SLC25A35 = 0.752 | 0.010; phosphorylated TP53BP1 = 0.776 | 0.002; UBAP2L = 0.835 | 0.002 **B.** Absolute quantification (copies per cell) of TAPAs at 72 hours, measured using SureQuant MHC. Asterisks indicate q < 0.05 by unpaired two-sided t-test. Average approximate copies/cell DMSO vs. TMZ: phosphorylated AMPD2 = 57 vs. 93, q = 0.010; CSF3 = 2,478 vs. 33,739, q = 0.004; GDF15 = 80 vs. 1,577, q = 0.0003; HAS1 = 678 vs. 1,589, q = 0.006; IFI30 = 16,960 vs. 33,042, q = 0.006; PARP1 = 262 vs. 501, q = 0.006; RETSAT = 349 vs. 601, q = 0.010; SLC25A35 = 56 vs. 133, q = 0.006; phosphorylated TP53BP1 = 71 vs. 134, q = 0.006; UBAP2L = 594 vs. 1,185, q = 0.008. **C.** Log₂FC (treatment/average 0.05% DMSO TMT signal) of candidate TAPAs at 72 hours following low and high doses of: TMZ (10 µM, 900 µM), BI 907828 (Brigimadlin) (10 nM, 1 µM), radiation (2.5 Gy, 9 Gy), and IFN-γ (10 ng/mL, 100 ng/mL). **D.** Abundance ranking of all identified MHC-I peptides based on precursor ion intensity from recurrent GBM patient tumors, with TMZ TAPAs highlighted. **E.** Correlation analysis between average log₂FC (TMZ vs. DMSO at 72 hours) of MHC-I peptides and their corresponding transcripts (DMSO R² = 0.06, TMZ R² = 0.08), proteins (DMSO R² = 0.02, TMZ R² = 0.02), and newly synthesized proteins (DMSO R² = 0.02, TMZ R² = 0.06). **F.** Comparative analysis of average log₂FC (TMZ/DMSO TPMs or TMT signal at each timepoint) for each TAPA’s mRNA, total protein, newly synthesized protein, and/or phosphosites. Asterisks indicate q < 0.05 (unpaired two-sided t-test, TMZ vs. DMSO).

We validated the identity and quantified the TMZ response of these 10 TAPAs using the SureQuant MHC method with stable isotope-labeled (SIL) versions of each peptide.^19^ Heavy-labeled TAPAs (100 fmol) were loaded onto recombinant HLA monomers and spiked into cell lysates to estimate absolute endogenous quantities using targeted, internal standard-triggered parallel reaction monitoring (IS-PRM). Endogenous TAPA levels ranged from hundreds to tens of thousands of copies per cell at 72 hours (Fig. 5B, Supplementary Fig. S5). The GDF15-derived peptide exhibited a striking ∼10-fold increase in TMZ-treated cells compared to DMSO, rising from baseline levels near ∼100 copies per cell, highlighting its potential as a treatment-inducible therapeutic target.

### TMZ TAPAs are also induced by other DNA-damaging therapies

To evaluate if these TAPAs were broadly induced by genotoxic stress, we treated U87-MG cells with additional DNA-damaging therapies, including high and low doses of TMZ, radiation, and the MDM2 inhibitor brigimadlin, which activates p53 signaling in patient-derived xenograft GBM mouse models,^72^ and quantified peptide abundance using discovery-mode, TMT-based mass spectrometry. IFN-γ was included as a control treatment, given its known enhancement of MHC-I presentation via the cGAS-STING pathway.^73^ Despite differing mechanisms of action, most TAPAs had increased expression following TMZ, radiation, and MDM2 inhibitor treatment, suggesting their potential roles in communicating DNA damage and p53 activation to the immune system (Fig. 5C). Interestingly, IFN-γ did not induce presentation of these peptides, except for GDF15 and PARP1. Peptides derived from RETSAT and phosphorylated TP53BP1 were consistently induced across all three genotoxic treatments. The CSF3 peptide was selectively upregulated by TMZ and radiation, suggesting a dependence on direct DNA damage signaling, whereas GDF15 was broadly induced across high-dose treatments, including IFN-γ, consistent with its role as a general stress-response marker.

### TMZ TAPAs in U87-MG cells are also detected in recurrent GBM patient tumors

To explore the clinical relevance of these TAPAs, we examined if they could be detected in human GBM tumor specimens. Using unlabeled discovery-mode MS, we analyzed two cohorts of recurrent tumors. The first comprised first-recurrence tumors from patients primarily treated with radiation and TMZ, some of whom had usually long survival, and the second consisted of patients with multiple recurrences and varied treatments (immunotherapy, gene therapy, chemotherapy, small molecule inhibitors) (Supplementary Table S2).

Several of the TMZ TAPAs were detectable in the patient tumors, albeit generally at mid-rank intensities, reflecting the vast diversity of self-peptides presented on MHC-I (Fig. 5D). In the first-recurrence cohort, UBAP2L, PARP1, and phosphorylated TP53BP1 peptides were identified in two unusually long-surviving patients (∼8-9.5 years post diagnosis). PARP1 and UBAP2L were also detected in moderately long-surviving patients (∼2-3.8 years): PARP1 in a patient who lived nearly 2 years, PARP1 and UBAP2L together in a patient who lived 3 years, and PARP1 again in a patient who survived ∼3.8 years with 4 recurrences. Additionally, IFI30 and UBAP2L peptides were detected in a patient who survived ∼3 years with 5 recurrences. While the absence of some TAPAs in other tumors may reflect differences in HLA alleles, other contributing factors could include the transient nature of MHC-I peptide presentation or immune escape mechanisms. Nevertheless, the detection of several TAPAs in recurrent GBM tumors supports their relevance as treatment-induced antigens that can be presented in patients.

### Changes in the immunopeptidome are not broadly correlated with the transcriptome, proteome, or translatome

To evaluate the relationship between MHC-I peptide presentation and internal cellular processes, we compared the change in expression induced by TMZ treatment of the immunopeptidome against corresponding changes in the transcriptome, proteome, and translatome in U87-MG cells. Across all samples, no significant global correlations were observed (Fig. 5E), consistent with previous reports.^11^ These findings highlight the challenge of predicting MHC-I peptide dynamics based solely on upstream gene or protein expression data.

Despite the lack of global correlation, several individual cases demonstrated coordinated changes across multiple layers of cellular regulation. The CSF3, GDF15, HAS1, IFI30, RETSAT, SLC25A35, and phosphorylated TP53BP1 peptides exhibited concordant upregulation at MHC-I peptide and other omics levels at 72 hours (Fig. 5F).

Conversely, other peptides displayed striking discordance between antigen presentation and upstream expression layers. For example, the MHC-I peptides derived from PARP1 and UBAP2L were significantly upregulated at 72 hours, despite both their transcripts and newly synthesized proteins being downregulated, suggesting regulation through alternative pathways such as preferential peptide processing or increased protein degradation. Similarly, the phosphorylated AMPD2-derived MHC-I peptide was significantly upregulated, as was intracellular phosphorylation at serine 114, while AMPD2 mRNA and translation were downregulated and total protein expression only marginally changed (Fig. 5F).

These findings illustrate that while some MHC-I peptides reflect altered transcript, protein, or protein synthesis levels, others are shaped by post-transcriptional or post-translational modifications, degradation dynamics, altered antigen processing, or binding affinities. This highlights the immunopeptidome as a distinct and informative readout of the cellular response to genotoxic stress, potentially revealing therapy-specific effects on immune peptide presentation.

## Discussion

Our systems-level analysis revealed that TMZ rapidly triggers cellular reprogramming in GBM cells, within a few days of exposure. This early response includes both the induction of mechanisms associated with longer-term resistance and a coordinated enhancement of MHC-I antigen presentation, particularly of previously uncharacterized TMZ-associated TAPAs. These findings suggest that standard treatment could heighten GBM immune visibility, a notable opportunity given GBM’s low tumor mutational burden (TMB) and fewer neoantigens compared with other cancers.^74^ By expanding the antigenic landscape, TMZ could be paired with immunotherapies such as cancer vaccines or BiTEs, both of which can access the brain but remain under-optimized for GBM.^14,75^ Our detected TAPAs included phosphorylated peptides, a class associated with increased MHC-I binding affinity and immunogenicity.^69,70^ Recent advances in phospho-MHC peptide detection methods may reveal additional TAPAs in GBM and other cancers.^76^ Another notable TAPA was one derived from GDF15, with strikingly consistent upregulation across transcriptional, translational, and immunopeptidomic layers. Given GDF15’s established role in tumor progression, this highlights a potential target for GBM. CRISPR-Cas9-mediated knockout of GDF15 has enhanced immunotherapy efficacy in syngeneic GBM mouse models,^77^ and anti-GDF15 antibodies have reversed immunotherapy resistance in clinical trials for non-small cell lung and urothelial cancers.^78^ However, targeting GDF15 is challenging due to its broader stress-induced expression in multiple tissues.^79^ Nevertheless, the identification of a GDF15-derived MHC-I peptide suggests a novel mechanism by which GDF15 may influence immune modulation in GBM and warrants further exploration.

TMZ-induced DNA damage may activate the cGAS-STING pathway, leading to the type I interferon responses we observed in the transcriptome, proteome, and at the cell surface. Several TAPAs were not induced by IFN-γ alone but required more direct DNA damage and p53 activation, suggesting the existence of treatment associated peptides distinct from those generated by classical inflammatory responses. We propose that DNA damage activates p53 and other stress pathways, slows down global translation, and reshapes the immunopeptidome. This parallels observations in melanoma, where disrupted translations leads to the presentation of immunogenic, targetable cancer antigens.^80^ Interestingly, some of the proteins that increased in new protein synthesis under TMZ are known to be translated through non-canonical mechanisms under stress. For example, TRIM22 represses cap-dependent translation by disrupting the eIF4F complex,^81^ and DDB2 contains an internal ribosome entry site in its 5’ UTR that allows cap-independent translation during stress.^82^ These non-canonical modes of translation may allow certain stress-adaptive proteins to continue being produced despite broader suppression of protein synthesis. Additionally, the increased translation of proteins involved in ubiquitination and protein degradation (CUL2, DDB2, TRIM22) may facilitate enhanced protein turnover and subsequent peptide presentation on MHC-I complexes and contribute to the increased immunopeptidome diversity generated by TMZ.

However, even if TMZ reliably induces TAPAs, these alone are unlikely to generate durable anti-tumor immunity, consistent with the limited benefit of TMZ monotherapy. Several factors may contribute to this, highlighting the limitations of this study: first, the immune system may not recognize these TAPAs, as they may not be efficiently cross presented by antigen presenting cells (APCs). We did not assess the immune relevance of the TMZ TAPAs and important next steps include evaluating the immunogenicity of these peptides, for example, by incubating TAPA-presenting GBM tumor cells with human T cells, as well as generating TAPA-targeting immunotherapies such as BiTEs or antibody-drug conjugates. Second, the notoriously immunosuppressive environment of GBM may prevent T cell infiltration or function,^14^ even in the presence of TAPAs. Cancer vaccines and BiTEs have struggled to advance beyond early phase clinical trials in GBM,^14,75^ and the tumor microenvironment, intratumoral heterogeneity, and immune context likely influence these outcomes. Expanding analyses to patient-derived tissues such as slice cultures or patient-derived cells, immune-competent mouse models, or 3D organoid systems will better recapitulate the tumor-immune interface. Third, our cell line, U87-MG, while well-characterized and practical for hypothesis generation and testing, does not represent the heterogeneity of GBM. Although we detected these TAPAs in 6 out of 15 recurrent tumors, which increased our confidence in their translational relevance, this was a small sample set that would benefit from the expanded model systems previously mentioned. Future studies across larger and more representative GBM cohorts will also be important to determine whether factors such as MGMT promoter methylation, a key clinical predictor of TMZ response, affect the prevalence or persistence of TAPAs, and to clarify TAPAs occurrence in patients with more typical disease courses.

Finally, these TAPAs may be transiently expressed. Our cell culture experiments captured the first 72 hours after TMZ exposure, though several were detected in patient tumors much longer after exposure. Determining whether these TAPAs persist, recur, or diminish over longer treatment intervals will be critical for translating these findings into viable therapies. Notably, several of the early stress-related responses we observed across multiple molecular layers are associated with therapeutic resistance. This suggests that resistance programs may emerge concurrently with TAPAs, underscoring the potential value of front-loading combinations with immunotherapies.

Overall, our findings show that within days of TMZ exposure, GBM cells undergo stress and antigenic remodeling, generating treatment associated immune peptides that could broaden the repertoire of targetable antigens in this otherwise low-TMB cancer. Though the immunogenicity and durability of these TAPAs require further exploration, they present a potentially exploitable window in which immunomodulatory effects of DNA damaging agents could be coupled with targeted immune activation. More broadly, this multi-omics approach enables exploration of the early immunologic consequences of genotoxic therapies in other cancers, where TAPAs may similarly create opportunities to enhance existing treatments.

## Acknowledgements

The authors thank the following members of the Forest White lab for feedback on methodology and technical support: Yufei Cui (immunopeptidomics), Daniel Rothenberg (translatomics), and Cameron Flower (bioinformatics). We also thank Stuart Levine of the MIT BioMicro Center for RNA sequencing support; Michele Griffin (flow cytometry), Alla Leshinsky and Richard Sciavoni (polymer synthesis), and Charlie Whittaker (bioinformatics) of the MIT Swanson Biotechnology Center for their assistance. Additional thanks to Alissa Caron at the Mayo Clinic and Joey Orpilla at UCLA for their assistance with recurrent GBM tumor sample transfers. This work was supported by NIH grants U54 CA283114, R01 CA080024, R21 ES036341, the MIT Center for Precision Cancer Medicine, NIH Training Grant in Environmental Toxicology T32-ES007020, the Superfund Research Program Grant P42 ES027707, and the following MIT Koch Institute student research funds: Charles S. Krakauer Fund, Hope Babette Tang (1983) Student Research Fund, Kristin R. Pressman & Jessica J. Pourian (2013) Fund, and Pearl Staller Graduate Student Fund. ChatGPT was used to assist with grammatical editing of this manuscript. All output was carefully reviewed, verified, and editing by the authors, who take full responsibility for the content.

