## Supplemental Figures for "Systems analysis uncovers early temozolomide responses and peptide antigens in glioblastoma"

Choe et al.

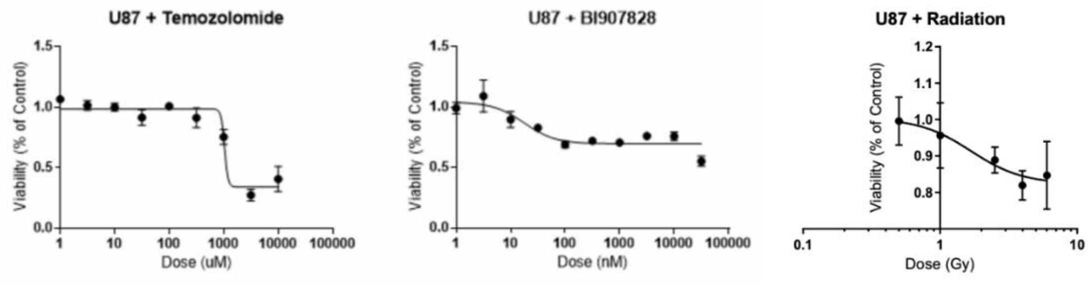


**Supplementary Figure S1.** Dose-response of U87-MG to TMZ, MDM2 inhibitor, and radiation (measured by CellTiterGlo).

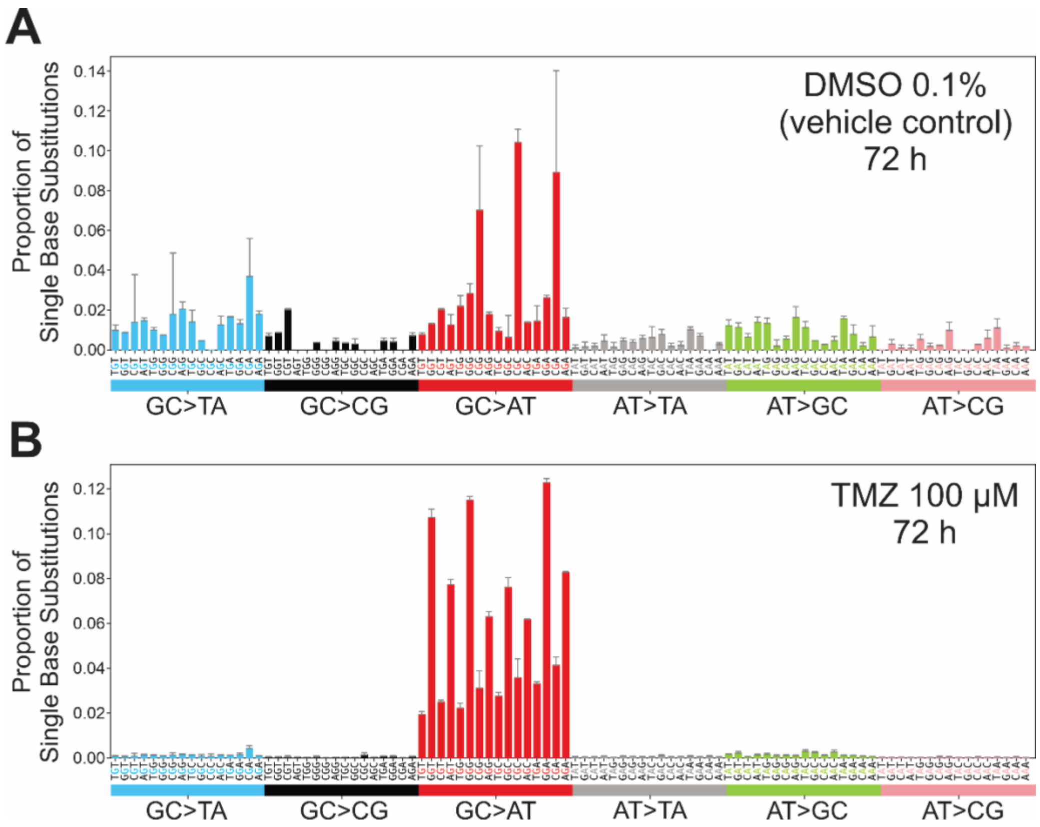


**Supplementary Figure S2.** Mutational spectrum of U87-MG cells at 72 hours post-treatment. Genomic DNA was isolated from U87-MG cells treated with DMSO (vehicle control) or 100 µM TMZ for 72 hours and analyzed by duplex sequencing. Single base substitution mutations were enumerated and plotted as a function of the trinucleotide sequence contexts in which they occur. Each data point reflects the average of 3 independent replicates, with the error bars denoting one standard deviation.

**A.** The mutational spectrum of DMSO control is typical of mutational spectra of cells growing in cell culture, with primarily GC→AT mutations in CpG sequence contexts (mutational signature SBS1) and GC→TA mutations reflecting 8-oxoguanine formation due to basal levels of oxidative stress.

**B.** The TMZ mutational spectrum features GC→AT mutations primarily in 5'-YGN-3' sequence contexts (mutational signature SBS11). This sawtooth pattern reflects the formation and miscoding properties of the $O^6$-methylguanine adduct, the key type of DNA damage driving the therapeutic effect of TMZ. The TMZ-induced mutation frequency in the U87-MG GBM cells, which reflects the upstream formation of O6-mG:T mismatches, is $3.32 \pm 0.05 \times 10^{-6}$, a 19-fold increase over the mutation frequency seen in the control samples ($1.7 \pm 0.3 \times 10^{-7}$). The TMZ mutational spectrum also provides information about the MGMT status of the cells. While O6-mG is the major mutagenic adduct generated by TMZ, a smaller amount of other DNA adducts is also formed; these minor adducts are responsible for mutations other than G→A, in particular AT→GC mutations. The extremely low (~2%) proportion of AT→GC mutations (green bars) observed here is indicative of cells that are MGMT deficient, which is consistent with prior knowledge regarding MGMT activity in U87-MG cells.

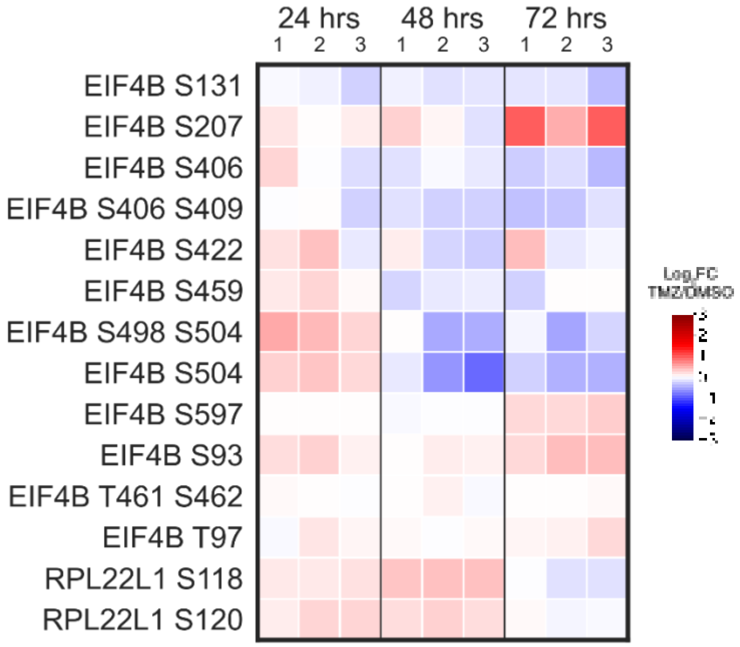


**Supplementary Figure S3.** Non-ATM/ATR phosphorylation of translation-related proteins. $Log_2FC$ (TMZ vs. DMSO) of TMT intensities are relative to the average TMT signal of corresponding DMSO-treated samples at each timepoint. Sites are global serine/threonine phosphorylation sites (not ATM/ATR substrate phosphorylation). All phosphosites shown had q-values > 0.05 (TMZ vs. DMSO, unpaired two-sided t-test, Benjamini-Hochberg correction) at 72 hours, except EIF4B S597, whose q-value at 72 hours = 0.044.

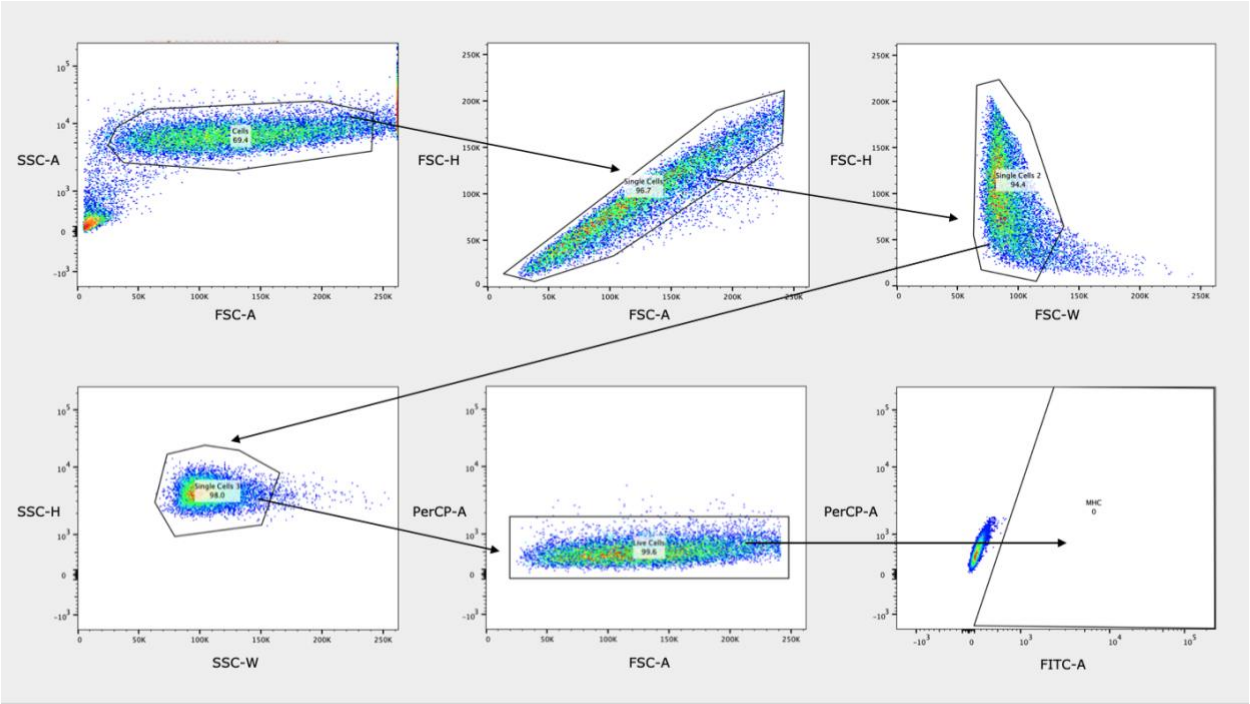


**Supplementary Figure S4.** Flow cytometry gating strategy. FSC-A vs. SSC-A was used to exclude debris. Doublets were removed sequentially using FSC-A vs. FSC-H, FSC-W vs. FSC-H, and SSC-W vs. SSC-H gates. Live cells were identified by DAPI exclusion using FSC-A vs. PerCP-A. Cells expressing MHC-I were gated using an Alexa Fluor 488-conjugated pan-HLA antibody, with FITC-A vs. PerCP-A plots and unstained controls (shown here) used to distinguish FITC+ from FITC- populations.

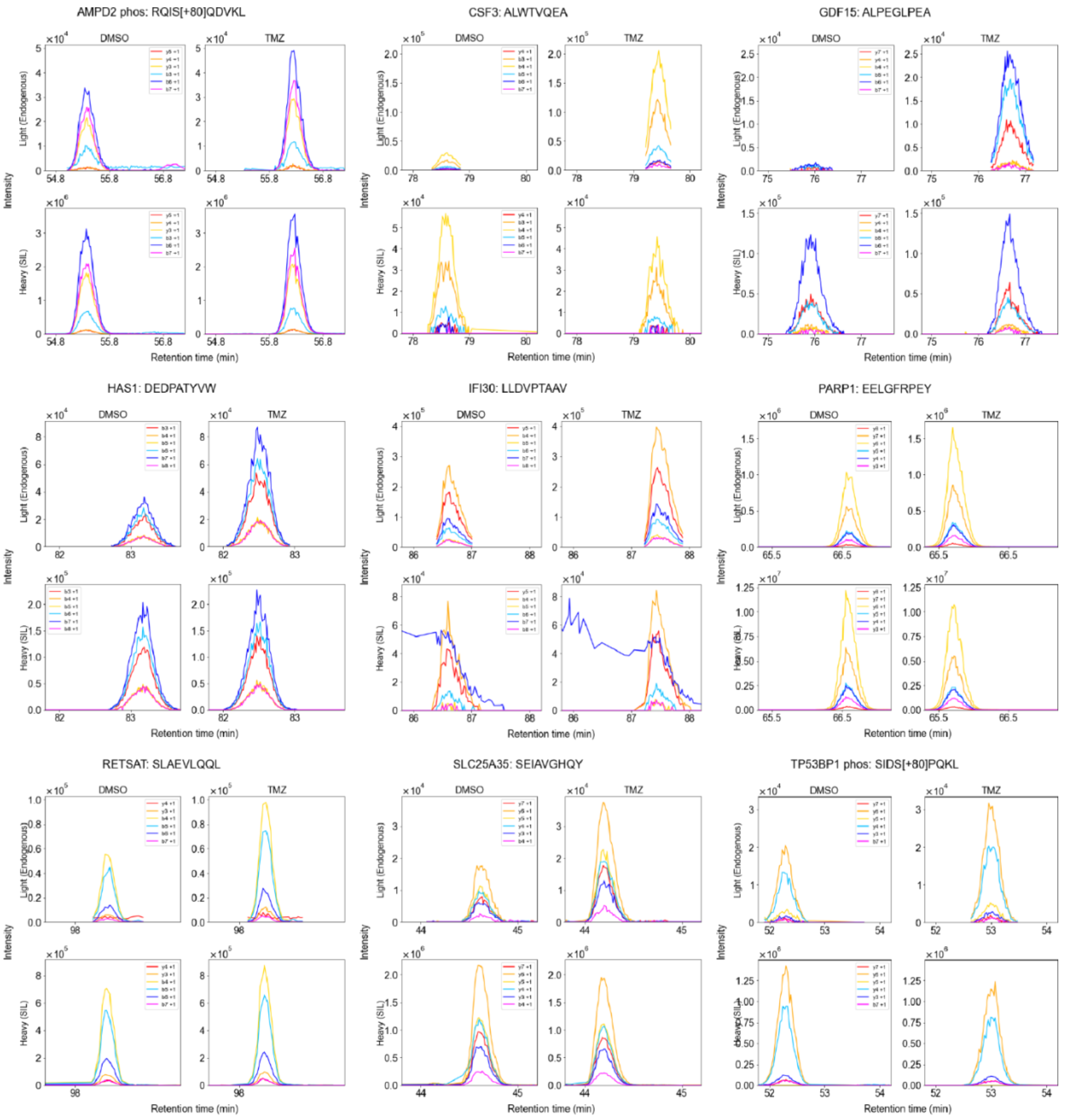


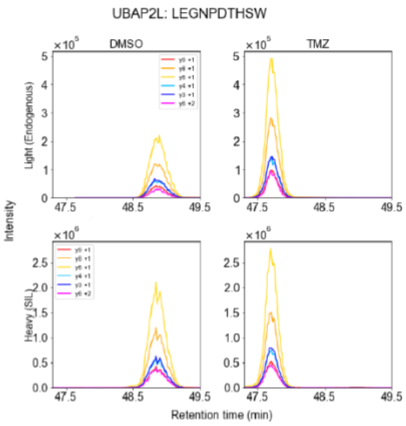


**Supplementary Figure S5.** Representative extracted ion chromatograms (XICs) for TMZ treatment-associated peptide antigens (TAPAs) and corresponding stable isotope-labeled peptides (SILs).

SUPPLEMENTARY METHODS

### Duplex sequencing

U87-MG cells were exposed to 100 µM TMZ dissolved in DMSO or 0.1% DMSO control for 72 hours in triplicate. The cells were washed with PBS, detached with TrypLE (Gibco), pelleted and stored at -80°C. Total genomic DNA was isolated using Monarch Spin gDNA Extraction Kit (New England Biolabs, Ipswitch, MA) using the manufacturer's protocol. One thousand nanograms of gDNA were enzymatically fragmented and used to construct duplex sequencing libraries using the Human Mutagenesis Kit (TwinStrand Biosciences, Seattle, WA) following the manufacturer's protocol. Following quality control determination of the DNA libraries, using FragmentAnalyzer and qPCR (BioMicroCenter, MIT), sequencing was performed on an Illumina NovaSeq 6000 using one lane of an S4 flowcell, utilizing a 150 bp paired-end kit (BioMicroCenter, MIT). The raw sequencing files were analyzed on the DNANexus computing cluster using the TwinStrand DuplexSeq Mutagenesis App v4.6.1. Data normalization and plotting was done using custom Python scripts as previously reported (Armijo et al., 2023; reference 22 in Main Text).

### LC-MS/MS analysis

All analyses used an Orbitrap Exploris™ 480 mass spectrometer (Thermo Fisher) with 2.5 kV spray voltage unless otherwise noted.

*Motif-specific phosphorylation*
Peptides were loaded onto an analytical chromatography column using a helium packing device. Columns were made in-house: fused silica capillary with inner diameter 50 µm and outer diameter 200 µm (Molex, catalog #1068150017) was pulled using a micropipette laser puller to create an integrated emitter tip with 1-2 µm inner diameter, then packed with 10 cm of 5 µm C18 beads (YMC, catalog # AQ12S05) and conditioned using a tryptic digest of bovine serum albumin. LC-MS/MS was performed using an Agilent 1100-Series chromatograph coupled to the mass spectrometer. Peptides were separated using 0.2 M acetic acid (solvent A) and 70% acetonitrile in 0.2 M acetic acid (solvent B) over the following 140-minute gradient: 0 min: 0% B, 10 min: 11% B, 115 min: 32% B, 125 min: 60% B, 130 min: 100% B, 133 min: 100% B, 140 min: 0% B at a flow rate of approximately 100 nL/min (200 µL/min through a flow splitter achieving a ~1:2000 split).

The mass spectrometer was operated in data-dependent acquisition (DDA) mode as follows: Full MS scan of precursor ions (m/z: 380–2,000, resolution: 60,000, AGC target: 300%) were followed by a series of MS/MS scans (monoisotopic peak determination, precursor intensity greater than 1e4, precursor charge state 2-6, isolation window: 0.4 m/z, HCD collision energy: 33%, resolution: 60,000, AGC target: standard, maximum injection time: 250 ms) for a total cycle duration of 3 seconds. Each precursor could be selected for MS/MS twice with a mass tolerance of +/- 10 ppm before being dynamically excluded for 45 seconds.

*Crude peptides*

Analysis of the supernatant of the motif-specific phosphopeptide IP was performed to correct for variations in peptide labeling across TMT channels. The supernatant was diluted 1:1000 in 0.1% acetic acid and approximately 30 ng (~3 µL) diluted supernatant was loaded onto a column (100 um ID x 10 cm) packed with 10 µm C18 beads (YMC) connected in series to an analytical chromatography column as described for the motif-specific phosphopeptides analysis. Liquid chromatography-tandem mass spectrometry was performed using an Agilent (1100-Series) chromatograph coupled to a Q Exactive Plus (Thermo Fisher) mass spectrometer. Peptides were separated using 0.2 M acetic acid (solvent A) and 70% acetonitrile in 0.2 M acetic acid (solvent B) over the following 75-minute gradient: 0 min: 0% B, 40 min: 30% B, 50 min: 60% B, 55 min: 100% B, 60 min: 100%, 65 min: 0% B, at a flow rate of approximately 100 nL/min (200 µL/min through a flow splitter achieving a ~1:2000 split).

The mass spectrometer was operated in DDA mode as follows: Full MS scan of precursor ions (m/z: 350–2,000, resolution: 70,000, AGC target: 3e6, maximum injection time: 50 ms) were followed by a series of MS/MS scans on the top 10 abundant ions (isolation window: 0.4 m/z, nominal collision energy: 33, resolution: 35,000, AGC target: 1e5, maximum injection time: 150 ms). Unassigned and +1 charge states were excluded and dynamic exclusion set to 30 seconds.

*Global phosphorylation (phospho-S/T) and total protein fractions*

Peptide fractions were loaded onto an analytical chromatography column like that used for motif-specific phosphopeptide analysis but packed with 3 µm instead of 5 µm C18 beads for improved chromatographic separation. Samples were directly loaded onto the analytical column using an UltiMate 3000 RSLC Nano LC system (Dionex) coupled to the mass spectrometer. A Nanospray Flex ion source (Thermo Fisher) was used with a column oven heater (Sonation) set to 50°C. Peptides were separated using 0.1% formic acid (solvent A) and 80% acetonitrile in 0.1% formic acid (buffer B) over the following gradient: 0 min: 3% B, 0.35 µL/min; 30 min: 3% B, 0.35 µL/min; 32 min: 6% B, 0.1 µL/min; 70 min: 19% B, 0.1 µL/min; 87 min: 29% B, 0.1 µL/min; 96 min: 41% B, 0.1 µL/min; 99 min: 97% B, 0.1 µL/min; 106 min: 97% B, 0.1 µL/min; 106 min: 3% B, 0.1 µL/min; 120 min: 3% B, 0.1µL/min.

The mass spectrometer was operated in DDA mode as follows: Full MS scan of precursor ions (m/z: 380–2,000, resolution: 60,000, AGC target: 300%) were followed by a series of MS/MS scans (precursor charge state 2-6, isolation window: 0.4 m/z, HCD collision energy: 33%, resolution: 60,000, normalized AGC target: 100%, maximum injection time: 150 ms) for a total cycle duration of 3 seconds. Each precursor could be selected for MS/MS 3 times with a mass tolerance of +/- 5 ppm before being dynamically excluded for 120 seconds.

*Translatome fractions*

Peptide fractions were loaded onto an in-house chromatography column like that used for motif-specific phosphopeptide analysis. Samples were directly loaded onto the analytical column using an UltiMate 3000 RSLC Nano LC system (Dionex) coupled to the mass spectrometer. A Nanospray Flex ion source (Thermo Fisher) was used with a column oven heater (Sonation) set to 50°C. Peptides were separated using 0.1% formic acid (solvent A) and 80% acetonitrile in 0.1% formic acid (buffer B) over the following gradient: 0 min: 3% B, 0.4 µL/min; 30 min: 3% B, 0.4 µL/min; 31 min: 3% B, 0.2 µL/min; 35 min: 14% B, 0.2 µL/min; 81 min: 42% B, 0.2

μL/min; 88 min: 60% B, 0.2 μL/min; 91 min: 96% B, 0.2 μL/min; 99 min: 96% B, 0.2 μL/min; 100 min: 3% B, 0.2 μL/min; 115 min: 3% B, 0.1μL/min.

The mass spectrometer was operated in DDA mode as follows: Full MS scan of precursor ions (m/z: 350–1,600, resolution: 60,000, AGC target: 300%) were followed by a series of MS/MS scans (precursor charge state 2-6, isolation window: 0.4 m/z, HCD collision energy: 33%, resolution: 45,000, normalized AGC target: 100%, maximum injection time: 150 ms) for a total cycle duration of 3 seconds. Each precursor could be selected for MS/MS twice with a mass tolerance of +/- 10 ppm before being dynamically excluded for 30 seconds.

*TMT-labeled immunopeptidome*
Peptide fractions were loaded onto an analytical chromatography column like that used for motif-specific phosphopeptide analysis but packed with 1.9 μm (ReproSil-Pur) instead of 5 μm C18 beads for improved chromatographic separation. Samples were directly loaded onto the analytical column using an UltiMate 3000 RSLC Nano LC system (Dionex) coupled to the mass spectrometer. A Nanospray Flex ion source (Thermo Fisher) was used with a column oven heater (Sonation) set to 50°C. Peptides were separated using 0.1% formic acid (solvent A) and 80% acetonitrile in 0.1% formic acid (buffer B) over the following gradient: 0 min: 3% B, 0.3 μL/min; 25 min: 3% B, 0.2 μL/min; 30 min: 6% B, 0.2 μL/min; 80 min: 25% B, 0.2 μL/min; 105 min: 35% B, 0.2 μL/min; 110 min: 55% B, 0.2 μL/min; 112 min: 97% B, 0.2 μL/min; 113 min: 97% B, 0.2 μL/min; 115 min: 3% B, 0.2 μL/min; 130 min: 3% B, 0.1μL/min.

The mass spectrometer was operated in DDA mode as follows: Full MS scan of precursor ions (m/z: 350–1,200, resolution: 60,000, AGC target: 300%) were followed by a series of MS/MS scans (precursor charge state 2-4, isolation window: 0.4 m/z, HCD collision energy: 33%, resolution: 60,000, AGC target: Standard, maximum injection time: Auto) for a total cycle duration of 3 seconds. Each precursor could be selected for MS/MS 3 times with a mass tolerance of +/- 5 ppm before being dynamically excluded for 120 seconds.

*Targeted, SureQuant quantitation of TAPAs*
Peptides were loaded onto an analytical chromatography column with the same parameters as the TMT-labeled immunopeptidome analysis over the following gradient: 0 min: 3% B, 0.4 μL/min; 20 min: 3% B, 0.4 μL/min; 25 min: 5% B, 0.2 μL/min; 90 min: 25% B, 0.2 μL/min; 100 min: 45% B, 0.2 μL/min; 103 min: 97% B, 0.2 μL/min; 104 min: 97% B, 0.2 μL/min; 105 min: 3% B, 0.2 μL/min; 115 min: 3% B, 0.2 μL/min.

DDA survey run: An initial DDA survey run of the 10 SILs of each TAPA was performed to determine the most intense product ions and optimal charge state of each SIL. SIL peptides were diluted in 0.1% formic acid and a mixture of 300 fmol-10 pmol of each peptide was injected in a single survey run, directly loaded onto an analytical chromatography column with the same parameters as the TMT-labeled immunopeptidome analysis over the following gradient: 0 min: 3% B, 0.4 μL/min; 20 min: 3% B, 0.4 μL/min; 25 min: 5% B, 0.2 μL/min; 90 min: 25% B, 0.2 μL/min; 112 min: 97% B, 0.2 μL/min; 115 min: 97% B, 0.2 μL/min; 116 min: 97% B, 0.2 μL/min; 117 min: 3% B, 0.2 μL/min; 130 min: 3% B, 0.2 μL/min.

The DDA analysis ran with an inclusion list of the precursor ions under +2, +3, and +4 charge states for each SIL peptide. The MS parameters were as follows: Full MS scan of precursor ions (m/z: 300–1500, resolution: 120,000, AGC target: 300%) were followed by a series of 20 MS/MS scans (isolation window: 1 m/z, HCD collision energy: 30%, resolution: 120,000, scan range: 350-1500 m/z, normalized AGC target: 1,000%, maximum injection time: 250 ms).

Targeted run: Peptides were directly loaded onto an analytical chromatography column with the same parameters and gradient as the DDA survey run. The custom SureQuant acquisition template available in Thermo Orbitrap Exploris Series 2.0 was used to build acquisition parameters for each m/z offset between SIL peptides and endogenous peptides. There were two nodes in total, one for peptides with a heavy glutamic acid or heavy lysine (both +6 m/z mass offset with +2 charge), and one for peptides with a heavy leucine (+7 m/z mass offset with +2 charge). The m/z of the peptides were specified in the "Targeted Mass Trigger" node, without an intensity threshold and with a mass tolerance of 5 ppm.

The MS parameters were as follows: Full MS scan of precursor ions (m/z: 380–1,200, resolution: 120,000, AGC target: 300%, maximum injection time: 50 ms) were followed by 30 MS/MS scans where heavy peptides within 5 ppm of the inclusion list items were isolated with an isolation window of 1 m/z, fragmented (HCD collision energy: 30%), with a scan range: 150-1,700, resolution: 15,000, normalized AGC target: 1,000%. A product ion trigger filter next performed pseudo-spectral matching, only triggering an MS/MS event of the endogenous, target peptide at the defined mass offset if n=3 of the top 6 product ions are detected from the defined list with a mass accuracy tolerance of 20 ppm. If triggered, the subsequent light peptide MS/MS scan had the same CE, scan range, and AGC target as the heavy trigger peptide, but with resolution 120,000.

*Unlabeled immunopeptidomes of GBM patient tumors*
Peptides were loaded onto an analytical chromatography column with the same parameters as the TMT-labeled immunopeptidome analysis.

Peptide fractions from each first-recurrence tumor (obtained from Mayo Clinic) were separated over the following gradient: 0 min: 3% B, 0.3 µL/min; 35 min: 3% B, 0.3 µL/min; 36 min: 6% B, 0.1 µL/min; 90 min: 30% B, 0.1 µL/min; 110 min: 45% B, 0.1 µL/min; 118 min: 55% B, 0.1 µL/min; 120 min: 97% B, 0.1 µL/min; 121 min: 97% B, 0.1 µL/min; 122 min: 3% B, 0.1 µL/min; 140 min: 3% B, 0.1 µL/min.

The mass spectrometer was operated in DDA mode as follows: Full MS scan of precursor ions (m/z: 350–1,200, resolution: 60,000, AGC target: 300%) were followed by a series of MS/MS scans (precursor charge state 2-4, m/z 350-1,200, isolation window: 0.4 m/z, HCD collision energy: 30%, resolution: 120,000, AGC target: standard, maximum injection time: 250 ms) for a total cycle duration of 3 seconds. Each precursor could be selected for MS/MS twice with a mass tolerance of +/- 10 ppm before being dynamically excluded for 30 seconds.

Peptides from each multiple-recurrence tumor (obtained from UCLA) were separated over the following gradient: 0 min: 3% B, 0.4 µL/min; 20 min: 3% B, 0.4 µL/min; 25 min: 6% B, 0.4 µL/min; 100 min: 25% B, 0.4 µL/min; 105 min: 45% B, 0.1 µL/min; 107 min: 100% B, 0.4 µL/min; 108 min: 100% B, 0.4 µL/min; 110 min: 2% B, 0.4 µL/min; 125 min: 2% B, 0.4 µL/min.

The mass spectrometer was operated in DDA mode as follows: Full MS scan of precursor ions (m/z: 350–1,200, resolution: 60,000, AGC target: 300%) were followed by a series of MS/MS scans with monosisotopic peak detection (precursor charge state 2-4, minimum intensity = 5,000, m/z 350-1,200, isolation window: 0.4 m/z, HCD collision energy: 30%, resolution: 45,000, AGC target: standard, maximum injection time: auto) for a total cycle duration of 3 seconds. Each precursor could be selected for MS/MS twice with a mass tolerance of +/- 10 ppm before being dynamically excluded for 30 seconds.

**Peptide identification and quantitation**

Raw files of mass spectra were processed using Proteome Discoverer version 3.0 (Thermo Fisher) and searched using Mascot version 2.4 (Matrix Science) with the canonical human proteome (SwissProt reviewed sequences, version 2023_01). For downstream analyses, corrected TMT intensities were summed for peptide identifications that were otherwise identical but differed only by methionine oxidation (commonly introduced during sample preparation) or by number of missed cleavages.

*Phosphoproteome*
Spectra were searched as follows: enzyme: trypsin, maximum missed cleavages: 1, precursor mass tolerance: 10, fragment mass tolerance: 20 mnu. Static modifications: cysteine carbamidomethylation. Dynamic modifications: TMT-labeled lysine, TMT-labeled peptide N-termini, methionine oxidation, tyrosine/serine/threonine phosphorylation. The Percolator node was used to calculate error probabilities and q-values for each peptide spectrum match (PSM). Phosphorylation site localization was performed using the ptmRS node.

TMT reporter ion intensities were extracted with an integration tolerance of 10 ppm and were isotope-corrected in Proteome Discoverer using according to the manufacturer-provided batch-specific isotopic impurities of each TMT channel.

PSMs were filtered for phosphorylation on SQ/TQ sites (for ATM/ATR substrates), SPXK/TPXK/SPXR/TPXR sites (for CDK substrates), or S/T sites (for global phosphorylation). PSMs were further filtered as follows: search engine rank = 1, isolation interference ≤ 30%, and at least one of the three options: ion score ≥ 25, q-value ≤ 0.01, and/or ion score ≥ 20 and q-value ≤ 0.05. PSMs with missing TMT values were discarded. TMT reporter ion intensities of PSMs with the same peptide sequence and phosphorylation site were summed. Reporter ion intensities were corrected for variations in sample labeling by using the median of peptide ratios in the crude peptide analysis for each channel relative to channel. Crude peptide spectra were searched and filtered with the same parameters as phospho-SQ/TQ spectra, but without phospho-tryrosine/serine/threonine modifications or ptmRS.

*Total protein and crude peptides*
Spectra were searched as follows: enzyme: trypsin, maximum missed cleavages: 2, precursor mass tolerance: 10, fragment mass tolerance: 20 mmu. Static modifications: cysteine carbamidomethylation, TMT-labeled lysine, TMT-labeled peptide N-termini. Dynamic modifications: methionine oxidation. The Percolator node was used to calculate error probabilities and q-values for each peptide spectrum match (PSM). PSMs were filtered using the

same quality control criteria—such as search engine rank and ion score cutoffs—applied to the phosphoproteomics data (excluding the initial phosphopeptide-specific filtering step).

*Translatome*
Spectra were searched as follows: enzyme: trypsin, maximum missed cleavages: 1, precursor mass tolerance: 10, fragment mass tolerance: 20 mmu. Static modifications: cysteine carbamidomethylation, TMT-labeled lysine, TMT-labeled peptide N-termini. Dynamic modifications: methionine oxidation, tyrosine/serine/threonine phosphorylation, 13C(6)15N(2) lysine, 13C(6)15N(4) arginine, TMT-labeled 13C(6)15N(2) lysine, methionine→AHA replacement, methionine→AHA with reduced azide replacement. The Percolator node was used to calculate error probabilities and q-values for each peptide spectrum match (PSM). Phosphorylation site localization was performed using the ptmRS node.

TMT reporter ion intensities were extracted with an integration tolerance of 10 ppm and were isotope-corrected in Proteome Discoverer using according to the manufacturer-provided batch-specific isotopic impurities of each TMT channel. PSMs were filtered as follows: search engine rank = 1, isolation interference ≤ 30%, and at least one of the three options: ion score ≥ 25, q-value ≤ 0.01, and/or ion score ≥ 20 and q-value ≤ 0.05. PSMs with missing TMT values were discarded. TMT reporter ion intensities of PSMs from the same protein were summed.

Variations in reporter ion intensities due to sample labeling were corrected using AHA-SHIP intensities. AHA-SHIPs were separately searched using a custom database with 13C(6)15N(1) isoleucine, 13C(6) arginine, TMT-labeled 13C(6) lysine, methionine→AHA replacement, methionine→AHA with reduced azide replacement, and methionine oxidation as dynamic modifications and cysteine carbamidomethylation, TMT-labeled lysine, and TMT-labeled peptide N-termini as static modifications. AHA-SHIP PSMs were filtered for search engine rank = 1, isolation interference ≤ 30%, and at least one of the three options: ion score ≥ 25, q-value ≤ 0.01, and/or ion score ≥ 20 and q-value ≤ 0.05. PSMs with missing TMT values were discarded. AHA-SHIP PSMs were filtered for ones with reporter ion intensities that fell within 10-fold of the interquartile range of all endogenous PSMs in the translatome run. Ratios of AHA-SHIP reporter ion intensities were calculated relative to the TMT126 channel, and the median of these ratios were used to correct each reporter ion intensity of the endogenous translatome PSM.

*TMT-labeled immunopeptidome*
Spectra were searched as follows: enzyme: trypsin, maximum missed cleavages: 1, precursor mass tolerance: 10, fragment mass tolerance: 20 mmu. Static modifications: TMT-labeled lysine, TMT-labeled peptide N-termini. Dynamic modifications: methionine oxidation, tyrosine/serine/threonine phosphorylation. The Percolator node was used to calculate error probabilities and q-values for each peptide spectrum match (PSM). Phosphorylation site localization was performed using the ptmRS node.

TMT reporter ion intensities were extracted with an integration tolerance of 10 ppm and were isotope-corrected in Proteome Discoverer using according to the manufacturer-provided batch-specific isotopic impurities of each TMT channel. PSMs between 8 and 15 amino acids long were filtered as follows: search engine rank = 1, isolation interference ≤ 30%, and at least one of the three options: ion score ≥ 25, q-value ≤ 0.01, and/or ion score ≥ 20 and q-value ≤ 0.05. PSMs with missing TMT values were discarded. TMT reporter ion intensities of PSMs with the same peptide sequence were summed.

Variations in reporter ion intensities due to sample labeling were corrected using hipMHC intensities. The hipMHCs were searched separately using a custom database with C-terminal amidation, 13C(6)15N(1) leucine, and methionine oxidation as dynamic modifications and TMT-labeled lysine and TMT-labeled peptide N-termini as static modifications. The hipMHC PSMs were filtered for search engine rank = 1, isolation interference ≤ 30%, and at least one of the three options: ion score ≥ 25, q-value ≤ 0.01, and/or ion score ≥ 20 and q-value ≤ 0.05. PSMs with missing TMT values were discarded. The hipMHC PSMs were filtered for ones with reporter ion intensities that fell within 10-fold of the interquartile range of all endogenous PSMs in the immunopeptidome run. Ratios of hipMHC reporter ion intensities were calculated relative to the TMT126 channel, and the median of these ratios were used to correct each reporter ion intensity of the endogenous immunopeptidome PSM.

*Targeted, SureQuant TAPAs quantitation*
Skyline Daily Build 24.1.1.202 was used to analyze SureQuant data of the 10 TMZ TAPAs. For each heavy peptide, the area under the curve (AUC) of the three most intense product ions was integrated, along with the AUC of the corresponding light, endogenous fragment ions in the same time frame. The ratio of the heavy:light product ions were calculated and averaged for each peptide, and this average ratio was multiplied by 100 fmol, the amount of SIL peptides spiked into each sample, to calculate the fmol of the endogenous peptides present in each sample.

*Human tumor immunopeptidomics*
Spectra were searched as follows: enzyme: trypsin, maximum missed cleavages: 1, precursor mass tolerance: 10, fragment mass tolerance: 20 mmu. Dynamic modifications: methionine oxidation, tyrosine/serine/threonine phosphorylation. The Percolator node was used to calculate error probabilities and q-values for each peptide spectrum match (PSM). The Minora Feature Detector node was used to estimate precursor abundances of each peptide spectrum match. PSMs between 8 and 15 amino acids long were filtered as follows: search engine rank = 1, isolation interference ≤ 30%, and at least one of the three options: ion score ≥ 25, q-value ≤ 0.01, and/or ion score ≥ 20 and q-value ≤ 0.05.
